# Seed bio-priming with phosphate solubilizing bacteria strains to improve rice (*Oryza sativa* L. var. FARO 44) growth under ferruginous ultisol conditions

**DOI:** 10.1101/2021.11.23.469751

**Authors:** Musa Saheed Ibrahim, Beckley Ikhajiagbe

## Abstract

The research investigated the possibility of phosphate solubilizing bacteria (PSB) with plant growth-promoting (PGP) capabilities to improve growth properties of rice plant under ferruginous ultisol (FU) condition through bio-priming strategy. The PSB with PGP properties used in this research were *Bacillus cereus* strain GGBSU-1, *Proteus mirabilis* strain TL14-1 and *Klebsiella variicola* strain AUH-KAM-9 that were previously isolated and characterized following the 16S rRNA gene sequencing. Biosafety analysis of the PSB isolates was conducted using blood agar. The rice seeds were then bio-primed with the PSBs at 3, 12 and 24 hours priming durations and then sown in a composite FU soil sample. Differences in germination bioassay involving SEM, morphology, physiology and biomass parameters were investigated for 15 weeks after bio-priming. The composite FU soil used in the study had high pH, low bioavailable phosphorus, low water holding capacity and high iron levels which has led to a low growth properties of rice seeds without bio-priming in FU soil. Germination parameters was better in seeds bio-primed with the PSBs, especially at 12h priming duration as against seeds without priming. SEM showed more bacterial colonization in the PSB bio-primed seeds. Seed bio-priming of rice seed with *Bacillus cereus* strain GGBSU-1, *Proteus mirabilis* strain TL14-1 and *Klebsiella variicola* strain AUH-KAM-9 under FU soil condition significantly improved seed microbiome, rhizocolonization and soil nutrient properties, thereby enhancing growth properties of the rice plant. This suggest the ability of PSB to solubilize and mineralize soil phosphate and improve its availability and soil property for optimum plant usage in phosphate stressed and iron toxic soils.

## Introduction

The application of synthetic fertilizers in improving soil nutrient has long been employed by conventional farmers. However, excessive application of these chemicals has led to accumulation of contaminants in agricultural soils (Lu et al., 2018) and consequently, a significant damage on important crops have been recorded (Musa and Ikhajiagbe, 2021; Herren et al., 2020). For example. in ferruginous soil which is characterized by iron-rich, acidic pH and phosphorus deficiency (Yu et al., 2016), application of phosphorus containing chemicals has further complicated the nutrient levels because, the phosphorus forms complexes with iron (Fe^3+^) in acidic soils, resulting in a ferrous phosphate (FePO) soil (Kumar et al., 2018; Satyaprakash et al., 2017). These processes still makes phosphorus not available for plants and heighten the soil acidity, which restrict the proper growth and yield of crops (Walpola and Yoon, 2012). Because of the high percentage (45.2 %) of Earth’s surface covered by the ferruginous soil and its high presence in some southern States of Nigeria such as Edo state (Ikhajiagbe et al., 2020; Doyou et al., 2017), it is important to develop an alternative strategy that can sustainably promote high crop yield under ferruginous soil condition.

Recently, the use of plant growth-promoting (PGP) bacteria has proven effective and sustainable in improving growth properties of plants (Musa, 2019; Suleman et al., 2018; Shen et al., 2016; Kuan et al., 2016). In particular, PGP bacteria has improved the availability of important soil nutrient through symbiotic N_2_-fixation and has improved the rate of secretion phtytohormones such as indole-acetic acid (IAA), cytokinins, and gibberellins (Shen et al., 2016). Backer et al. (2018) have reported an increase in physiological parameters of sweet pepper (*Capsicum annum*) as response to inoculation with PGP bacteria. Improved root length, surface area and general growth of Chick pea (*Cicer* arietinum) and Pigeon pea (*Cajanus cajan*) under zinc and copper concertation with the application PGP bacteria (species of *Enterobacter*) was also observed by (Ikhajiagbe and Musa, 2020a; Rizwan et al., 2018; Gopalakrishnan et al., 2016). However, in the case of ferruginous soil with high iron levels and low phosphorus, application of PGP bacteria alone may not be able to yield the required results because the PGP bacteria may not withstand the acidity and iron toxicity present in the soil (Zhao, 2014). For this reason, previous research have suggested the inoculation of phosphate solubilizing bacteria (PSB) for effective chelation of soil phosphorus to a bioavailable phosphorus for plant use (Wang et al., 2014). Several strategies for inoculation of bacteria to bring about improved growth properties have been explained (Rizwan et al., 2018; Gopalakrishnan et al., 2016). Previous researchers have attributed the success of PGP bacteria to the techniques used in inoculating the bacteria to the host plant (Mahmood et al., 2016; Gopalakrishnan et al., 2016). Foliar spray, rhizo-inoculation; which involve the introduction of microorganism into the root region of plants to stimulate growth have been previously tested by (Ikhajiagbe and Edokpolor, 2019). However, these techniques have certain limitations when it comes to efficiency in seed emergence and germination studies as described by Mahmood et al. (2016). While there are several other strategies of PGP bacteria inoculation into the host plant, seed bio-priming have been reported as one of the most effective ways to obtain a seed surface bacterial inoculation.

Bio-priming can be seen as a technique involving soaking of seeds using living bacterial inoculum for a particular duration to allow the bacterial imbibition into the seed (Ashraf and Foolad, 2005). This process initiates some physiological responses in seeds, which brings about fast colonization and acclimatization under controllable condition (Mahmood et al., 2016). According to Taylor and Harman (1990), bio-priming gives a sustainable condition for bacteria to continue multiplying rapidly and proliferate in the spermosphere, leading to enhancement in seed viability germination, germination speed, seed vigor, germination time, as well as improved growth and yield of plants (Ikhajigbe and Musa, 2020a; Rajendra et al., 2016). Bio-primed seeds have been documented as efficient in improving germinability and morphological performance of seeds under stressed environmental conditions (Mirshekari et al. 2012). Considering the effectiveness of bio-priming as a means of PGP bacteria inoculation, this research aims to investigate the possibilities of bio-priming rice (*Oryza sativa* L.) seeds with PGP bacteria having PSB capabilities to improve growth properties of rice plant under ferruginous condition.

The PGP bacteria having PSB capabilities used in this study were *Bacillus cereus* strain GGBSU-1, *Klebsiella variicola* strain AUH-KAM-9 and *Proteus mirabilis strain* TL14-1 which were previously isolated and identified by Musa and Ikhajiagbe (2020a) following the 16s rRNA, from ferruginous and humus soils in Benin City, Nigeria. The PSB isolates have proven efficient in solubilizing insoluble phosphate in soils (Musa and Ikhajiagbe, 2020a; Addendum 1 and 2) and plant growth-promoting capabilities by releasing IAA and siderophores (Musa and Ikhajiagb, 2020a; Addendum 3 and 4). Furthermore, these isolates have significantly improved germination and yield parameters in rice *in vitro* (Musa and Ikhajiagbe, 2021). The current study will bring about another sustainable strategy of improve agricultural productivity and food security under ferruginous ultisol condition.

## Materials and Methods

### Experimental soil preparation

The experiment was carried out at the experimental garden of the Department of Biology and Forensic Science, Admiralty University of Nigeria, Delta State Nigeria. Ferruginous soils that was previously obtained by Musa and Ikhajiagbe (2020b) from six locations around Benin City, Edo State of Nigeria were pooled to obtain a composite sample. The composite ferruginous soil was made in 12 sections prepared in experimental bowls (60 x 25 cm). Each section was prepared in five replicates.

### Soil physiochemical parameter

The ferruginous soil samples were air-dried at temperature of 22-25°C and then analyzed for soil organic matter levels (SOM), soil available phosphorus, cation exchange capacity (CEC), pH of the soil, total nitrogen, organic carbon (OC), exchangeable acidity (EA), available potassium, available micronutrients such as sodium (Na) and Aluminum (Al), electrical conductivity, soil texture class and maximum water holding capacity following Musa and Ikhajiagbe (2020a). The iron levels of the soil was analyzed following the method of Cheng et al. (2013) by using concentrated perchloric acid to digest the soil sample and subjecting it to titration with versanate solution as in Ikhajiagbe et al. (2019).

### Bacterial species

Three phosphate solubilizing bacteria species (*Bacillus cereus* strain GGBSU-1, *Klebsiella variicola* strain AUH-KAM-9 and *Proteus mirabilis* strain TL14-1) that were previously isolated from ferruginous ultisol and humus soil in an earlier study in Benin City by Musa and Ikhajiagbe (2021) were prepared in stock cultures for this study. The bacteria species were previously identified using molecular tool of 16S rRNA after biochemical test involving catalase, indole, citrate, nitrogen fixing activity and bromotyhmol blue test and pH tolerance level test with HCL following (Mondala *et al*., 2016). PGP capabilities of the isolates were determined by IAA and siderophores production following Gupta *et al*. (2012b) and Balkar, (2013) respectively and reported in Musa and Ikhajiagbe 2021. Meanwhile, the phosphate solubilizing capabilities were analyzed using pikovskaya’s media as reported in Musa and Ikhajiagbe (2021).

### Preparation of inoculum

The pure PSB having PGP traits (*Bacillus cereus* strain GGBSU-1, *Klebsiella variicola* strain AUH-KAM-9 and *Proteus mirabilis* strain TL14-1) were prepared by streaking on to agar plates and incubated at 28°C for 48 hours. After 48 hours growth, the isolates were inoculated in Nutrient broth and then prepared into 0.5 McFarland Standard with Cat. No (TM50) to standardize the approximate number of bacteria in the suspension. Following this process 500 mL of each bacteria isolate was prepared to obtain an average microbial suspension of 1.5×10^8^ cfu/mL.

### Biosafety analysis of the isolated PSBs

Since some strains of bacteria may be harmful to Humans (Muhamad *et al*., 2020), biosafety analysis of the isolated PSBs were carried out by using a blood agar medium in order to investigate their ability to grow in blood and such give information on its possible effect on Humans (Russell *et al*., 2006). The three PSB isolates were poured on an already prepared blood agar plate of 5% blood. The setup was incubated at 40°C for two days. Haemolysis of red blood cells was assessed by the formation of a clear zone around colonies (β-haemolysis), or greenish colouration (α-haemolysis), while no clear zone indicates γ-haemolysis.

### Rice seed bio-priming

Rice variety (FARO 44) seeds that was previously obtained from Center for Dryland Agriculture, Bayero University, Kano was used in this study. The seeds were tested for viability following (AOSA, 2000) and surface sterilized with 70% ethanol following Sameer and Nabeel (2016). The sterilized seeds were allowed to dry at room temperature for 3 hours and then sown onto 45 pre-sterilized petri dishes (9c diameter) at the rate of 10 seeds per petri dish that was lined with Whatman filter paper. 10 mL of the prepared 500mL bacteria inoculum of each bacteria was introduced into each petri dish and studied for 3, 12 and 24 hours to allow bacteria colonization. The control was made to be just 10 mL of distilled water. The set up was covered at a temperature between 25 and 26.6°C as described by Etesami et al. (2014). The three bacteria were coded as (A= *Bacillus cereus* strain GGBSU-1, B= *Proteus mirabilis* strain TL14-1 and C= *Klebsiella variicola* strain AUH-KAM-9), while the control containing 10mL of distilled water was coded with FD in ferruginous ultisol.

### Sowing of rice seeds

Three different bio-priming setup were transplanted after 3 hours, 12 hours and 24 hours into the already prepared experimental bowls containing the assayed ferruginous soils (Table 1). Each bacteria-primed seeds were transplanted into the 12 ferruginous soil sections at the rate of 10 seeds per pot. The set up was made in an open (no shade) environment and as such relied basically on rainfall. However, the soil moisture was always maintained periodically following the methods of USDA (2010). This set up was weeded at every two days maintained for 15 weeks after priming (WAP).

**Table 1:** Physical and chemical parameters of the experimental non-ferruginous (control) and ferruginous soil.

| Parameters | Ferruginous soil |
| --- | --- |
| Available phosphorus (mg/kg) | 8.01 ± 0.04 |
| Electric conductivity (μS/cm) | 301.09 ± 1.22 |
| pH | 5.01 ± 0.21 |
| Total organic carbon (%) | 0.41 ± 0.11 |
| Soil organic matter (%) | 7.31 ± 0.56 |
| Total Nitrogen (%) | 0.62 ± 0.01 |
| Exchangeable acidity (meq/100g) | 0.16 ± 0.13 |
| Cation exchange capacity (cmol/kg) | 1.70 ± 0.02 |
| Textural class | Loamy-sandy |
| Clay (%) | 10.92 ± 1.42 |
| Silt (%) | 8.72 ± 2.76 |
| Sand (%) | 95.10 ± 0.09 |
| Fe (mg/kg) | 200.67 ± 2.44 |
| Water holding capacity (%) | 68.89 ± 0.12 |
| Available potassium (mg/kg) | 0.02 ± 0.08 |
| Mg<sup>2+</sup> (meq/100g) | 4.21 ± 1.12 |
| Na<sup>+</sup> (meq/100g ) | 3.09 ± 0.29 |
| Al (meq/100g) | 6.23 ± 1.22 |

Fe = iron, Al= aluminum, Na= sodium, Mg= magnesium.

### Rice seed germination bioassay

Germination parameters are very effective in determining the influence of bacterial priming on seeds. Germination parameters were analyzed in two ways. First was determining the time at which the first coleoptile (primary leaf) appeared. This was studied *in vitro* at different bio-priming periods and observed by monitoring the setup after every 6 hours at 25°C in a dark room of the Department of Plant Biology and Biotechnology, University of Benin Postgraduate Research laboratory, following (ISTA, 2005).

The second was germination percentage which was studied *in vivo* after rice seeds have been transplanted into the soils. Germination was taken as the period at which the first cotyledon protrude visibly from the soil and calculated following (ISTA, 2005) as:

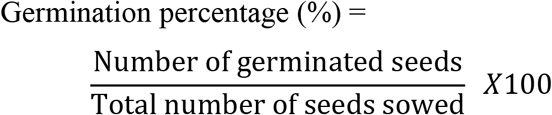

Bacterial colonization capabilities of the phosphate solubilizing bacteria (PSB) having plant growth promoting (PGP) traits (*Bacillus cereus* strain GGBSU-1, *Klebsiella variicola* strain AUH-KAM-9 and *Proteus mirabilis* strain TL14-1) on rice seed at 24hours after bio-priming was observed via electron microscopy. Endosperm sample (1 cm) were prepared aseptically and placed into vials for each treatment. The electron microscopy was designed following Muhamad et al. (2020) and viewed using scanning electron microscopy (SEM) Jeol JSM-6400.

### Morphological properties

Morphological parameters that are related to growth and yield of rice were investigated. Fresh shoot length, length of internode and length of the first leaf were calculated in (cm) by using a transparent ruler that was mounted on a white calibrated paper throughout the study. To obtain the dry shoot length (cm), seedlings were air dried for 24 hours and then measured from day 3 to 16 weeks after bio-priming. The length of internodes (cm) was measured from the coleoptile to the first node using a sample of 5 best seedlings from all treatments weekly. The average time taken for the formation of one leaf was determined as time per leaf formation (T/LF) following Ikhajiagbe et al. (2021) with slight modification as:

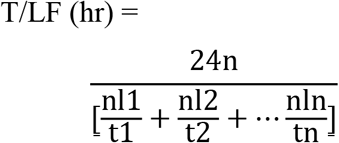

Where:

T/LF = time taken for the formation of one leaf
nl = number of leaves per unit time
n= number of periods
t = time during which parameter was measured

### Physiological parameters

To analyze the influence of bacteria bio-prime on the growth and yield of the rice plant, physiological parameters were investigated. Total soluble sugar (TSS) of fresh leaves were estimated from week 5 to week 14 with 2 weeks interval by oven drying the tallest leaf at 70°C for 24h as described by Nelson (1944) with slight modification by Sankar and Selvaraju (2015). Growth enzymes such as alpha amylase (AA) of the growing plant extract was determined at similar days as TSS by DNS method of Lowry *et al*. (1951) at a pH of 7.5. Terpenoid and lycopene of fresh leaves were determined at similar days as AA following the method of Moran (1982). Chlorophyll content index (CCI) of old and fresh leaves were determined using a non-destructive method employing Apogee chlorophyll concentration meter. The CCI of the old and new leaf were measured as average of the mesocotyl, mid seedling and top seedling at similar days as AA. Chlorophyll *a* and *b* contents were investigated according to methods described by Arnon *et al* (1949); Maxwell and Johnson (2000).

### Biomass parameters

Leaf area (cm^2^) was calculated using an android application (Leaf-IT) following Julian *et al*. (2017) at week 5 to week 14 with 2 weeks intervals. Number of leaves were measured by counting at every week till harvest. Leaf tip chlorosis and necrosis was measured as the percentage of the total number of leaves produced by plants that showed significant level of chlorosis and necrosis. This was pre-decided following reports from Ikhajiagbe *et al*. (2017) from week 3 to week 14, with 2 weeks interval. Weight of fresh leaf (g) was measured using analytical weighing balance at similar days as AA.

### Root growth parameters

Fresh root length were calculated in (cm) by using a transparent ruler that was mounted on a white calibrated paper throughout the study. To obtain the dry root length (cm), seedlings were air dried for 24 hours and the root were measured from day 3 to 16 weeks after bio-priming. Meanwhile, number of secondary roots were carefully observed and counted daily.

### Rice harvest properties

Other rice yield properties such as time at which maturity was achieved, number of tillers, number of reproductive tillers, number of panicle, straw yield, number of seeds per panicle, weight of rice panicle, weight of peduncle without rice, weight of 100 grains and plant tissue water were calculated at harvest day.

### Statistical analysis

Data obtained from the analysis were presented in means and standard errors of five replicates. Data were analyzed following two-way analysis of variance using GENSTAT (8th edition). Where significant p-values were obtained, differences between means were separated using Student Newman Keuls Test (Alika, 2006). The ferruginous soil used in the experiment was homogenized

## Results

### Soil physicochemical properties

Table 1 showed the physicochemical properties of the ferruginous soil used in this study. The result revealed that ferruginous soil is acidic with 5.01 pH and had a total soil organic carbon content of 7.31%. The iron content is high (200.67 mg/kg), while the available phosphorus level is low (8.01 mg/kg). The textural class was observed to be loamy-sandy with a poor water holding capacity (86.89%).

### Biosafety assay of the isolated PSBs

Results for PSB biosafety using blood agar showed that the three isolated PSBs did not exhibit any haemolytic activity, because no detectable clear zone (γ-haemolysis) was observed (Fig. 1).

**Fig. 1:**
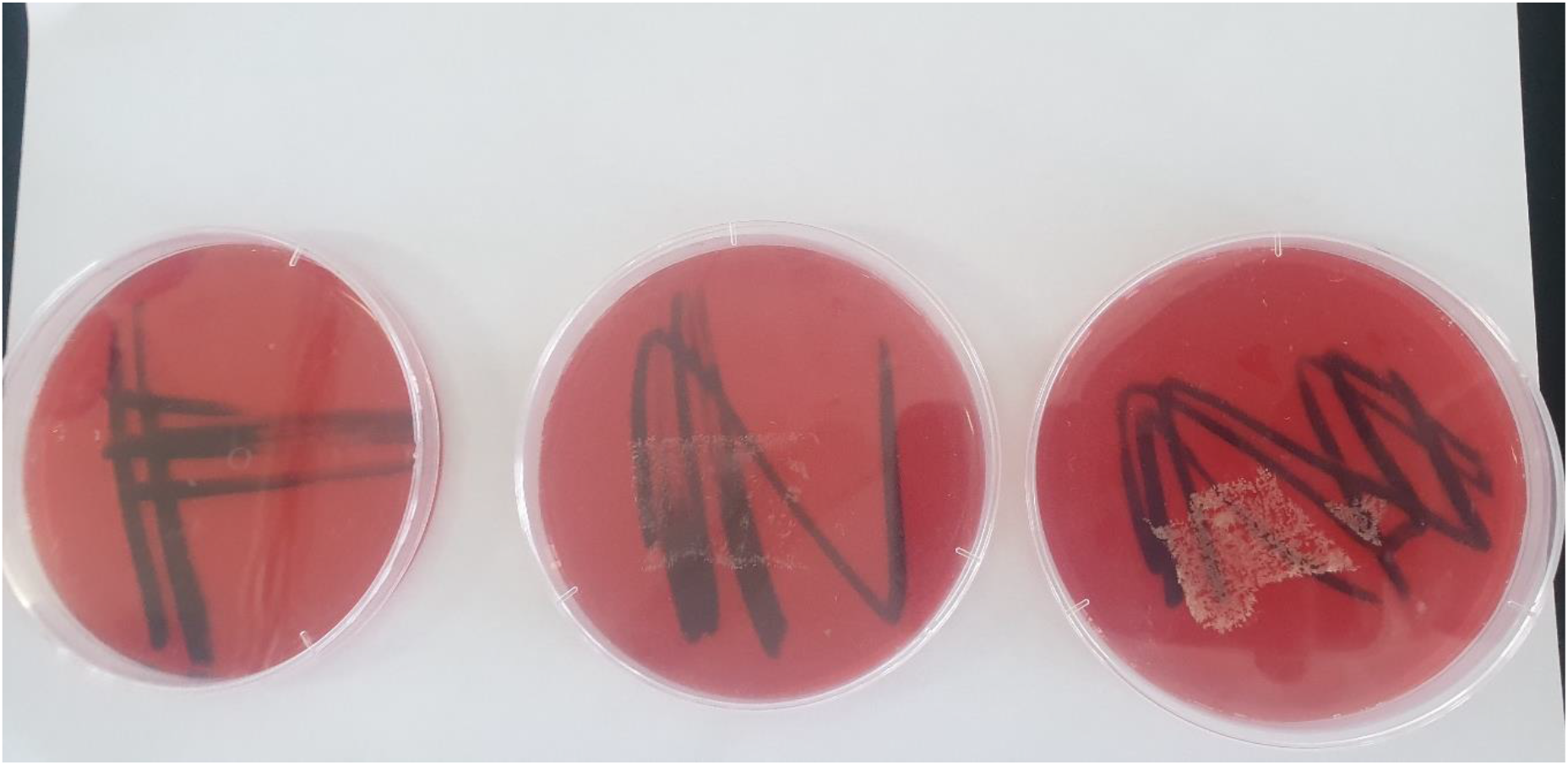
PSB isolates showing the (γ-haemolysis) with no clear zone. A= *Bacillus cereus strain GGBSU-1*. B = *Proteus mirabilis* strain TL14-1. C = *Klebsiella variicola* strain AUH-KAM-9.

### Germination assay and seed colonization of the three PSBs

Germination bioassay was carried out to investigate the influence of the three PSBs on germination parameters of rice seeds at the three different bio-priming durations. It was observed that the coleoptile did not appear in all the setup bio-primed for 3 hours. However, the first coleoptile appeared at 12 hours in the BPS (FA), followed by the PPS (FB) at 18 hours, while the KPS (FC) appeared at the 24^th^ hour (Table 2). No sign of coleoptile appearance was observed in the ferruginous soil without PSB inoculation (FD) throughout the 24 hours bio-priming experiment (Fig. 2). After transplanting, the BPS transplanted after 12 and 24 hours bio-priming (FA12 and FA24) was observed to be the first to achieve full germination (100%) at 2 days after transplanting (DAT), followed by the PPS (FB12) at 4 DAT (Table 2). Significant delay (p>0.5) was observed in the germination percentage of FD in all assayed days. Generally, the germination results indicated the highest positive influence of the BPS compared to PPS and KPS, while the FD showed the lowest influence (15 %).

**Table 2.** Germination parameters of rice seeds after 24hours bio-priming with three PSB isolates

| Samples | APC (Hrs) | Germination (%) on |  |  |
| --- | --- | --- | --- | --- |
|  |  | 2<sup>st</sup> DAT | 4<sup>nd</sup> DAT | 6<sup>rd</sup> DAT |
| FA3 | No Appearance | 82±1.3<sup>a</sup> | 92±1.1<sup>a</sup> | 100±2.1<sup>a</sup> |
| FB3 | No Appearance | 80±1.5<sup>a</sup> | 91±0.2<sup>a</sup> | 100±2.4<sup>a</sup> |
| FC3 | No Appearance | 80±2.3<sup>a</sup> | 89±1.9<sup>a</sup> | 100±2.6<sup>a</sup> |
| FD3 | No Appearance | 20±1.1<sup>c</sup> | 60±1.3<sup>b</sup> | 63±5.1<sup>b</sup> |
| FA12 | 12±1.5<sup>f</sup> | 100±2.3<sup>a</sup> | 100±2.4<sup>a</sup> | 100±0.4<sup>a</sup> |
| FB12 | No Appearance | 95±2.1<sup>a</sup> | 100±2.1<sup>a</sup> | 100±1.2<sup>a</sup> |
| FC12 | No Appearance | 95±3.3<sup>b</sup> | 98±0.9<sup>b</sup> | 100±1.8<sup>a</sup> |
| FD12 | No Appearance | 34±2.1<sup>c</sup> | 63±1.2<sup>c</sup> | 70±2.1<sup>b</sup> |
| FA24 | 12±1.5<sup>f</sup> | 100±1.0<sup>a</sup> | 100±2.1<sup>a</sup> | 100±5.7<sup>a</sup> |
| FB24 | 18±2.5<sup>f</sup> | 98±0.1<sup>a</sup> | 98±2.1<sup>a</sup> | 100±2.5<sup>a</sup> |
| FC24 | 24±2.5<sup>f</sup> | 90±3.4<sup>b</sup> | 96±3.0<sup>a</sup> | 100±2.5<sup>a</sup> |
| FD24 | No Appearance | 34±1.2<sup>c</sup> | 63±1.1<sup>b</sup> | 80±2.5<sup>b</sup> |

Results with similar alphabetic superscripts on same column did not differ from each other ( $p>0.05$ ). Results were in mean and standard error of five replicates. DAT=Day after transplant, APC= Appearance of coleoptile. FA3 = Ferruginous soil bio-primed with *Bacillus* sp for 3hours duration, FB3 = Ferruginous soil bio-primed with *Proteus* sp for 3hours duration, FC3 = Ferruginous soil bio-primed with *Klebsiella* sp. For 3hours duration. FD= Ferruginous soil without bio-priming. The number after each sample signifies duration of priming.

**Fig. 2:**
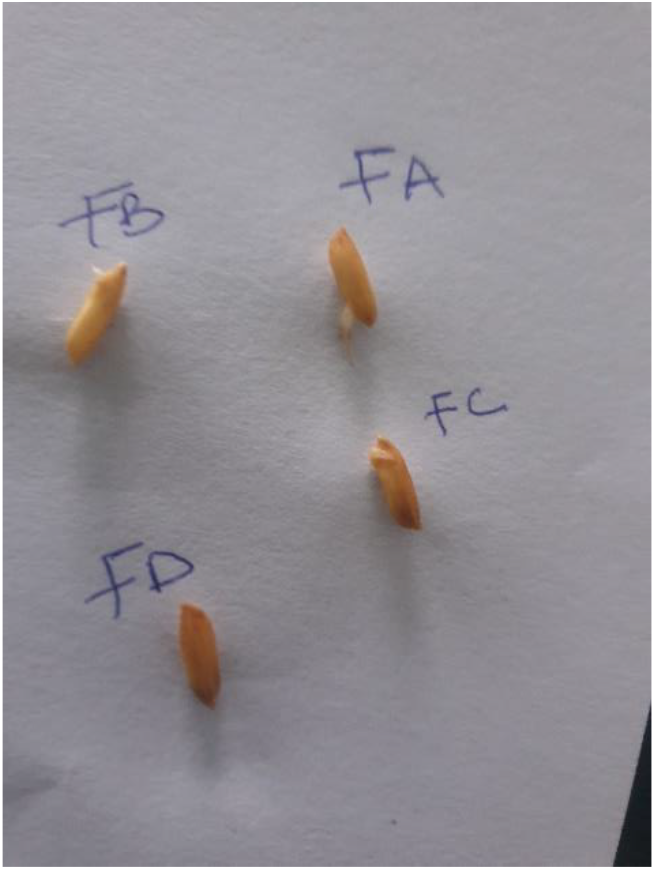
Appearance of Coleoptile at 24 hours after bio-priming.

To confirm the effective colonization of the three PSBs on rice seeds *in vitro*, the tin endosperm section of the rice seed bio-primed with the three PSBs and the control at 24 hours after bio-priming were viewed using SEM. Images of endosperm sections (Fig. 3) viewed under 3000× magnification showed that the three PSBs (*Bacillus cereus strain GGBSU-1*, *Proteus mirabilis* strain TL14-1. Respectively and *Klebsiella variicola* strain AUH-KAM-9) has successfully colonized and proliferate the endosperm at different cell proliferation levels. The BPS showed highest bacterial population, while the control showed no bacterial population.

**Fig. 3:**
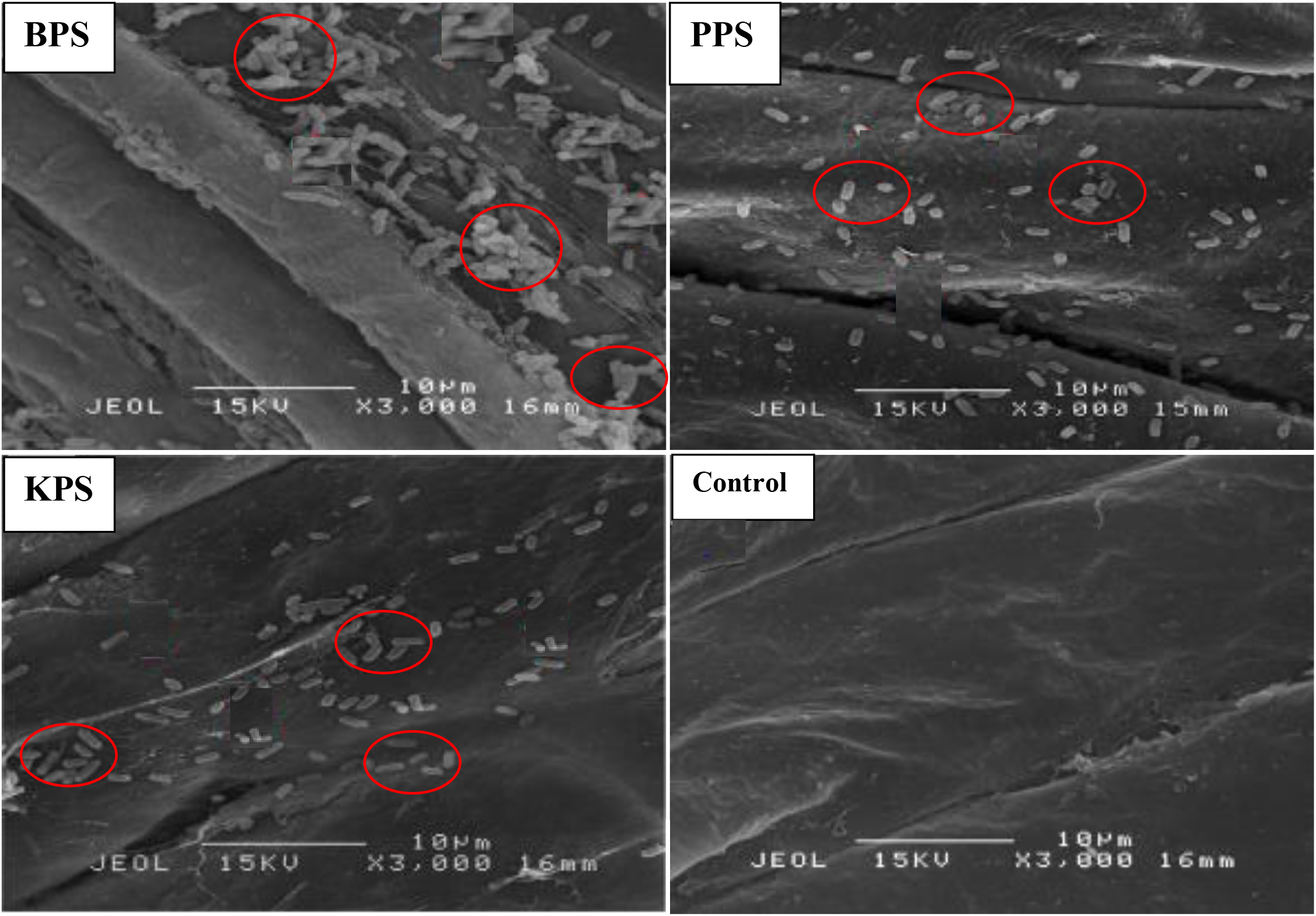
Images of rice endosperm sections showing in situ colonization and acclimatization of PSBs (indicated by oval) under SEM. BPS= *Bacillus cereus strain GGBSU-1* primed seed. PPS = *Proteus mirabilis* strain TL14-1 primed seed. KPS = *Klebsiella variicola* strain AUH-KAM-9 primed seed. Control = Rice seeds without PSB priming.

### Effect of bio-primed PSB on rice morphological parameters

Since the SEM imaging (Fig. 3) indicated bacterial colonization as a results of bio-priming, a seedling pot experiment was carried out to evaluate the possibilities of the PSBs to solubilize phosphate in ferruginous soil to improve rice plant morphology. Differences (Fig. 4, 5, 6, table 3 and Addendum 5) were observed in rice seedling morphological parameters between the PSB bio-primed seeds in ferruginous soils and the control (ferruginous soil without PSB priming). A significant increase (p<0.05) was observed in the fresh shoot length (Figure 4) of all PSB bio-primed seeds as compared to the ferruginous soil without priming. The FA12 was observed to show highest growth performance in all the assayed days, while FD24 was observed to show the least morphological properties. Dry shoot length, length of first leaf and length of internodes were all improved in the PSB bio-primed set up as against the ferruginous soil without priming. At 8WAT, delayed shoot length and length of internodes were observed. Furthermore, table 3 showed that rice seeds bio-primed with the three PSB bacteria took significantly lower period (33% low) for single leaf formation as compared to the non-primed seeds.

**Fig. 4:**
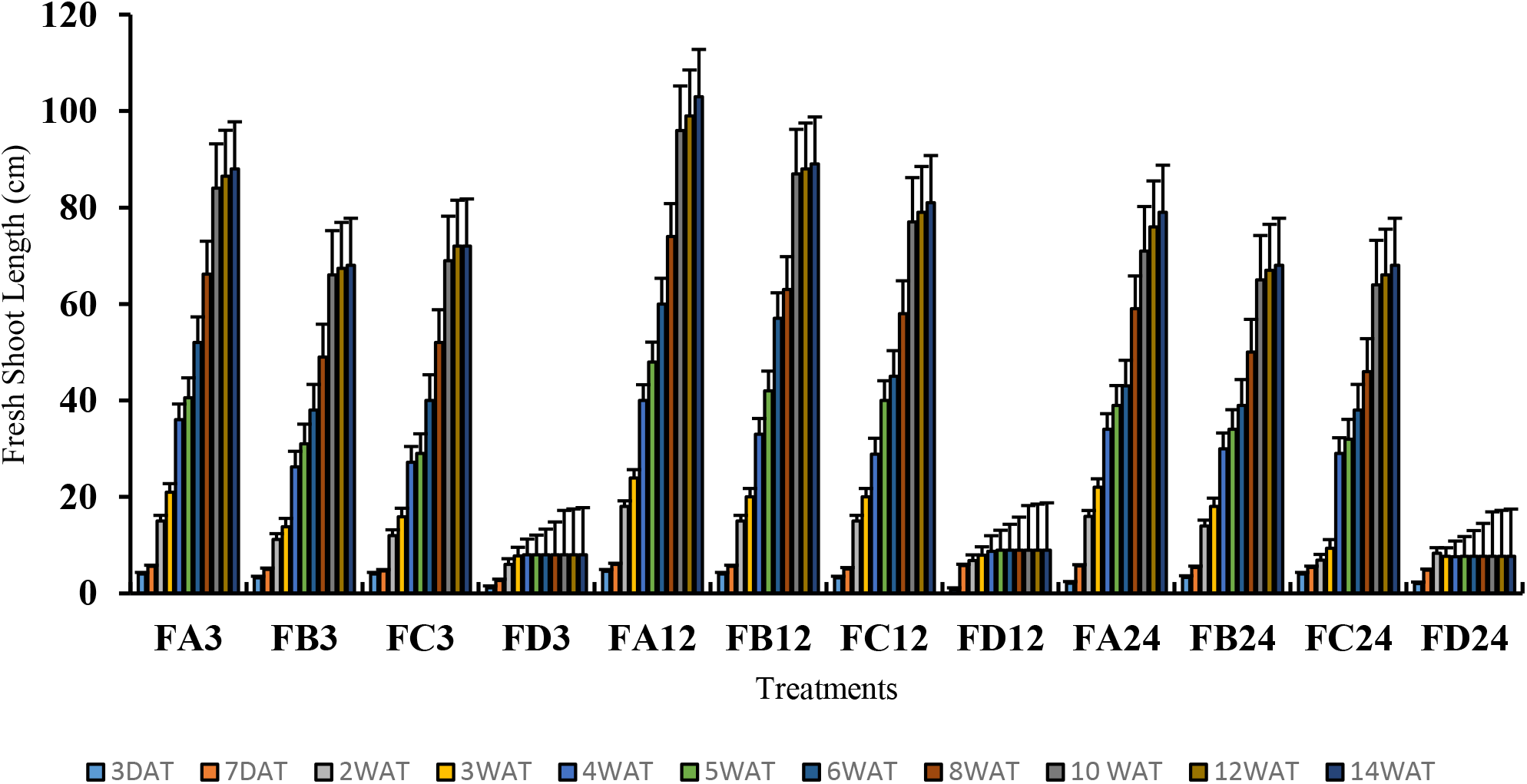
Fresh shoot length of rice from 3days after priming to 14 weeks after priming.

**Fig. 5:**
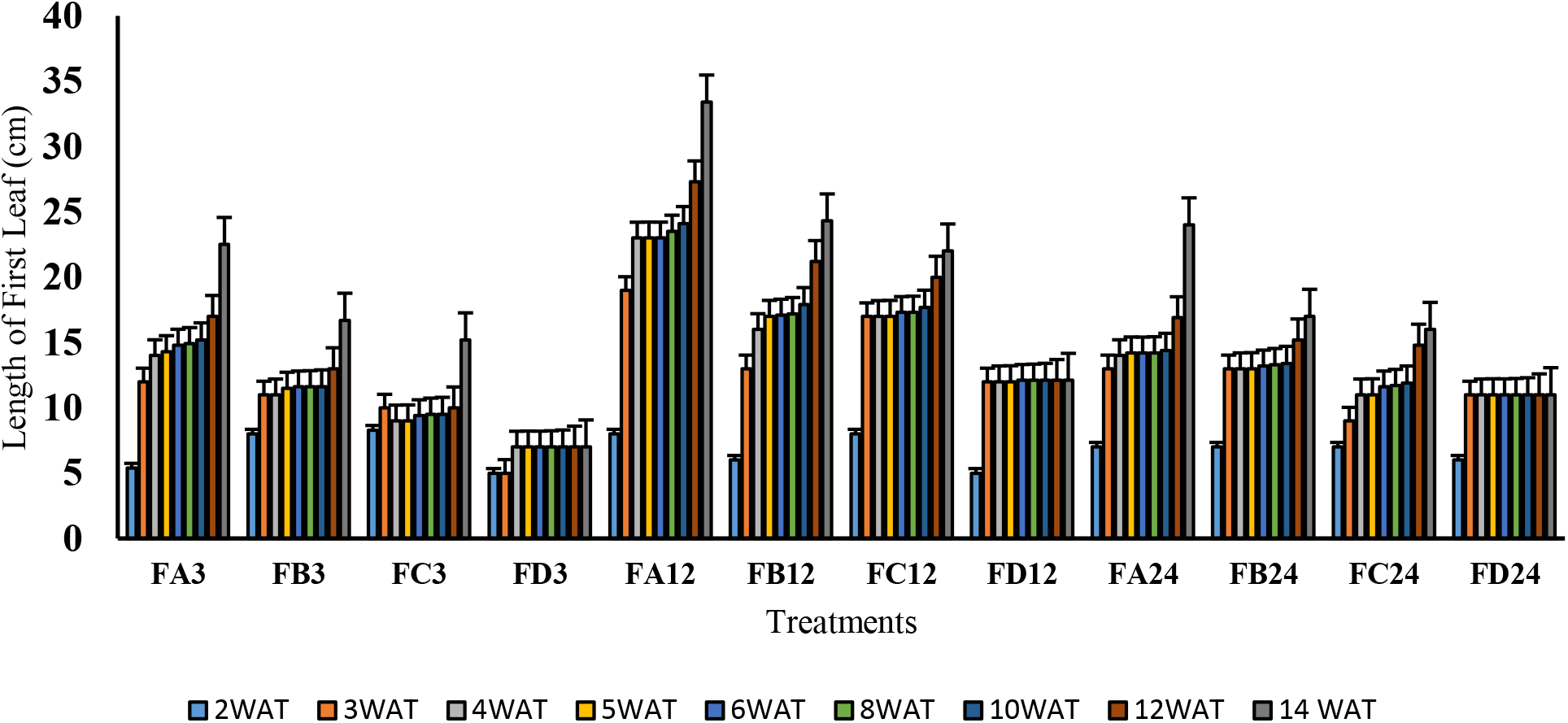
Length of First Leaf of rice from 3days after bio-priming to 14 weeks after bio-priming.

**Fig. 6:**
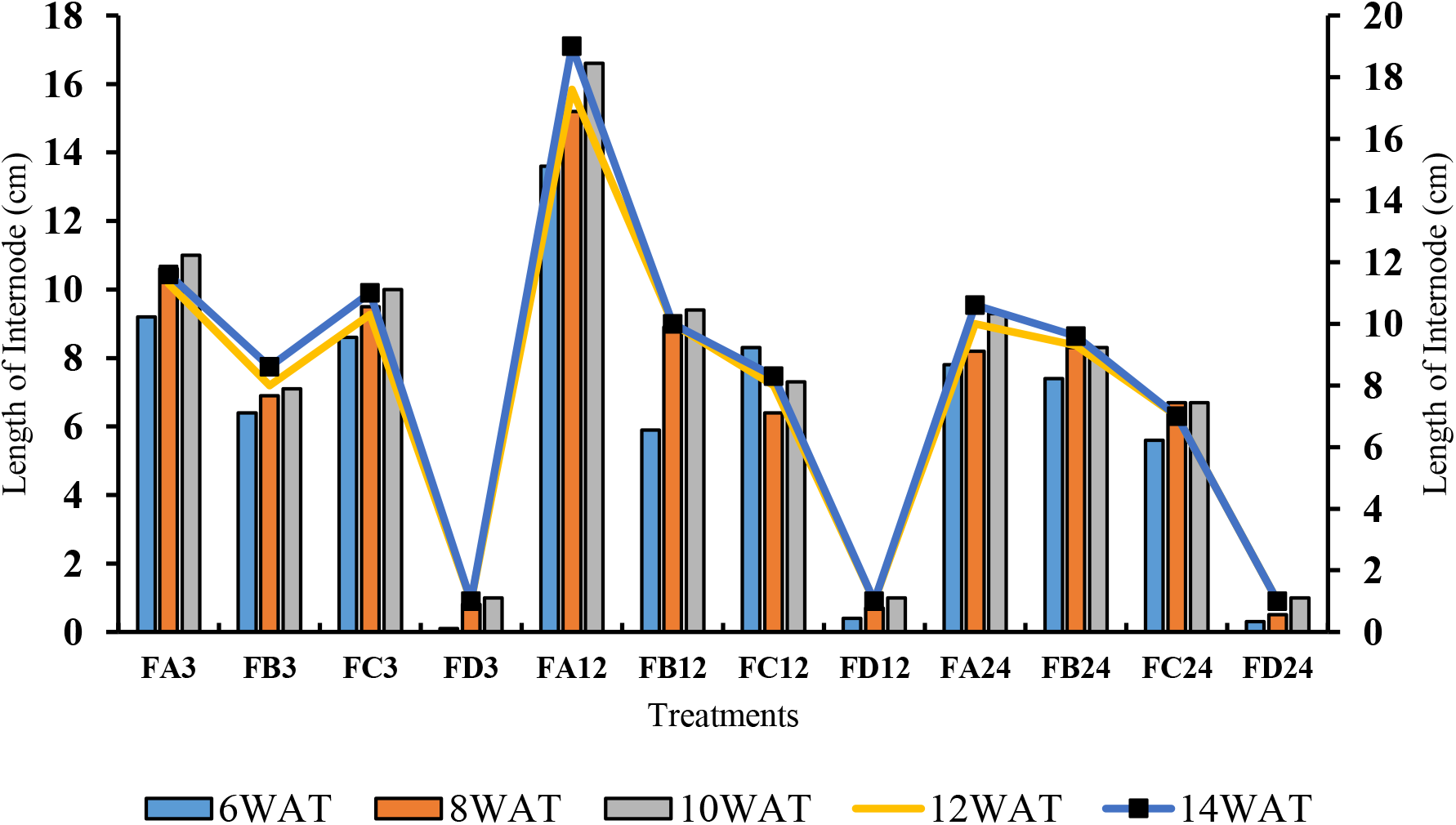
Length of Internode of rice from 3days after bio-priming to 14 weeks after bio-priming.

**Table 3.**
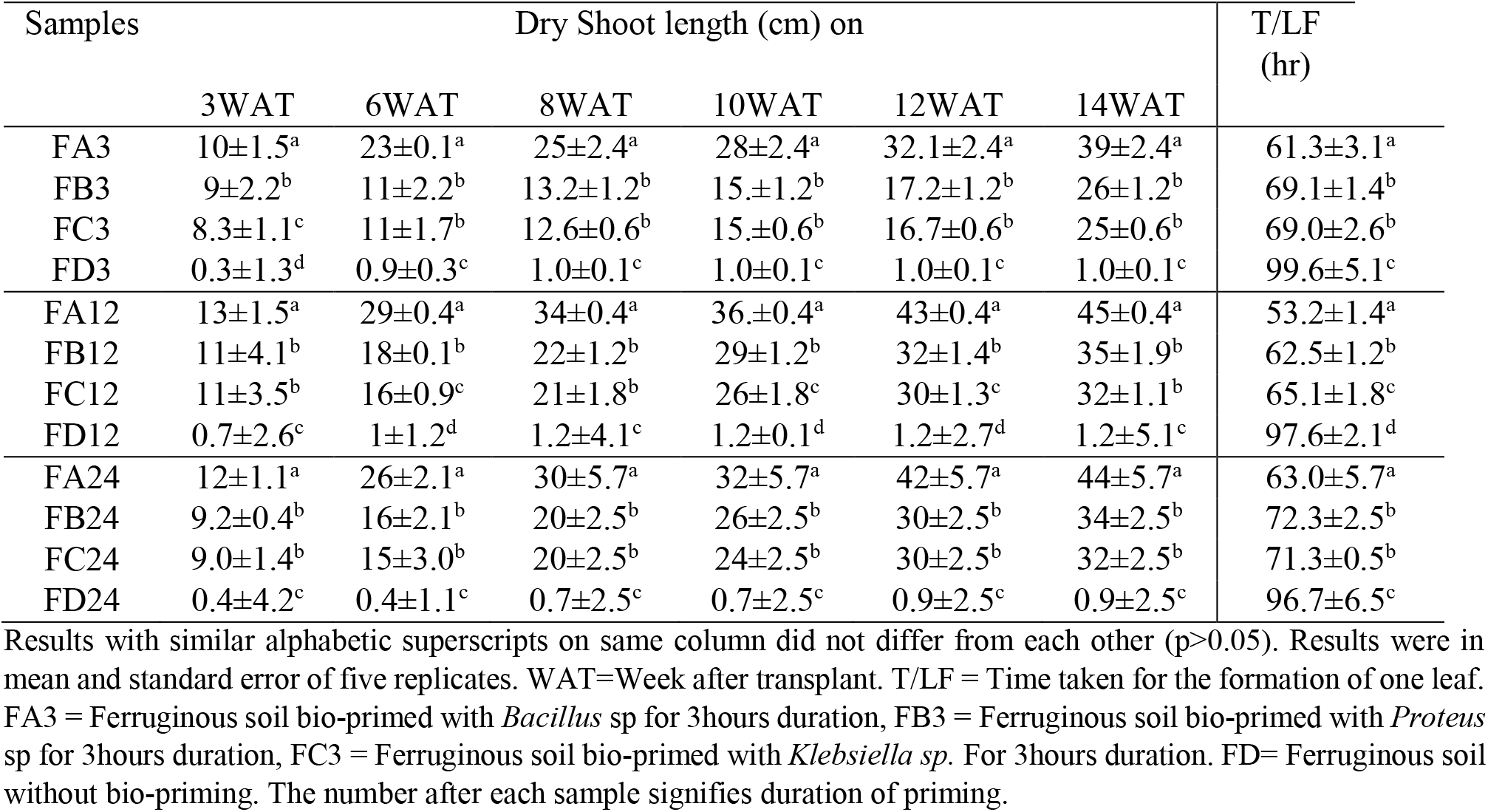
Response of dry shoot length and time taken for rice leaf formation to PSB bio-priming.

### Effect of bio-primed PSB on rice physiological parameters

Since terpenoid signals biotic and abiotic stress in plants, terpenoid levels was assayed in order to observe the influence of ferruginous soil on rice plant and the role of bio-priming. The result (Fig. 7) showed increased terpenoid levels in all the rice plant without bio-priming at all durations, while the rice plant that was bio-primed with PSBs showed lower levels. Significant increase in lycopene levels were observed in the bio-primed rice plant as against the non-primed set up (Fig. 8). However, the FA12 was observed to show highest levels. Significant differences (Fig. 9, 10 and 11) were observed in total soluble sugar (TSS), alpha amylase (AA) and chlorophyll content index (CCI) between the bio-primed rice plant and the non-primed plant. The BPS was observed to show highest (45%) TSS and AA levels as compared to KPS and PPS. Generally, old and new leaf CCI followed similar trend as in TSS and AA. However, a slight difference were observed between old leaf CCI and new leaf CCI, with the old leaf having more photosynthetic properties than the new leaf (Figure 11).

**Fig. 7:**
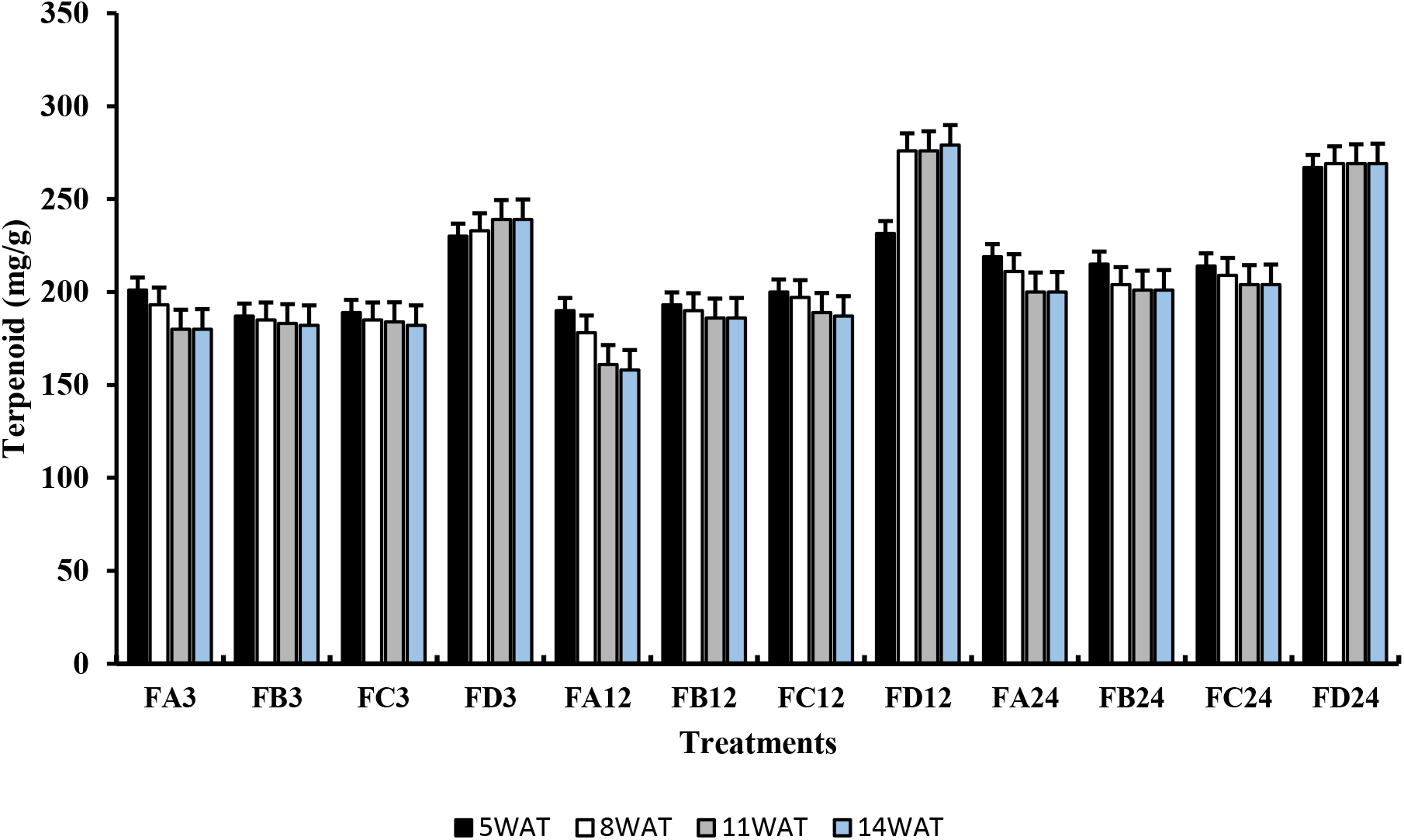
Terpenoid levels of rice plant from 3days after priming to 14 weeks after priming.

**Fig. 8:**
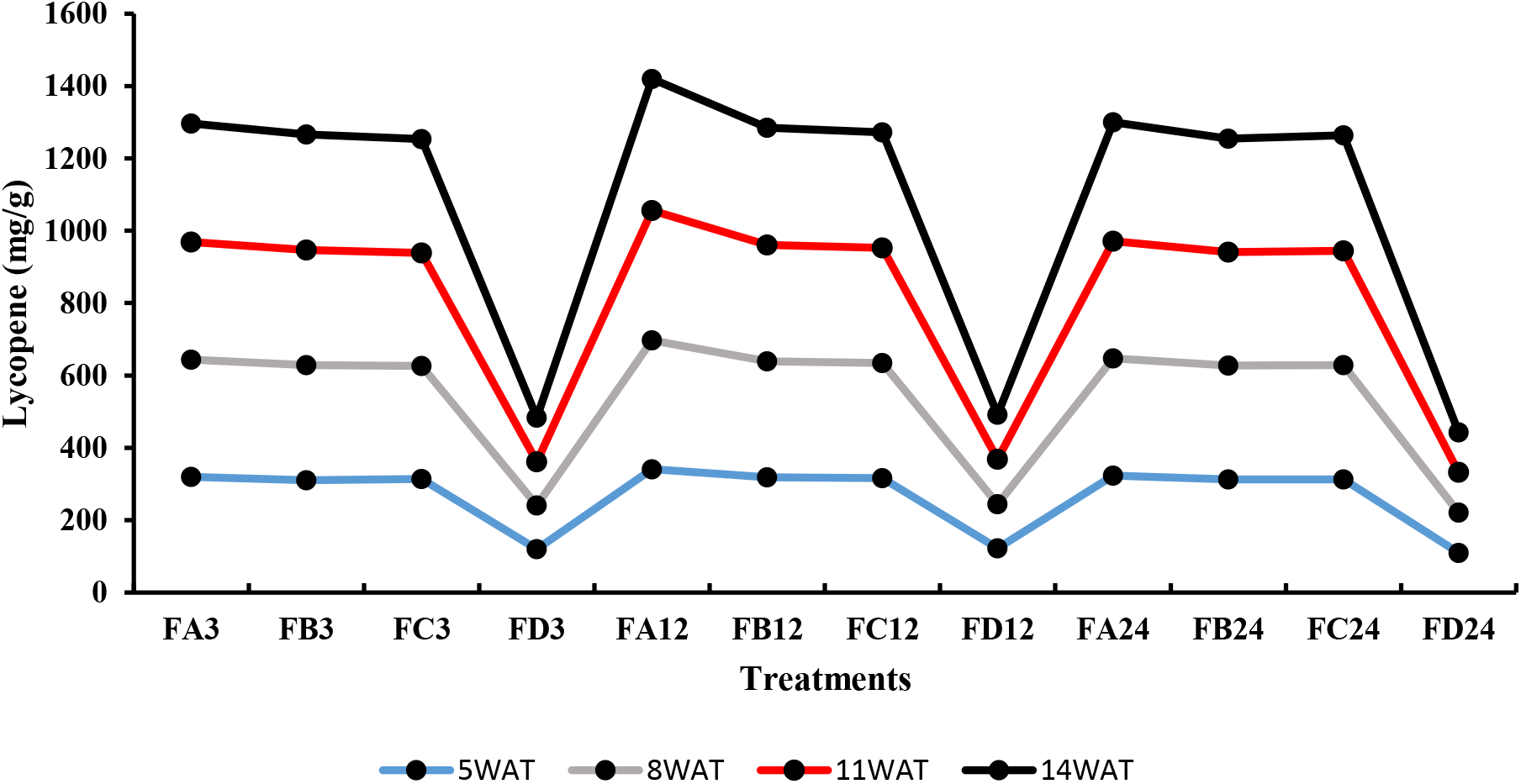
Lycopene levels of rice plant from 3days after priming to 14 weeks after priming.

**Fig. 9:**
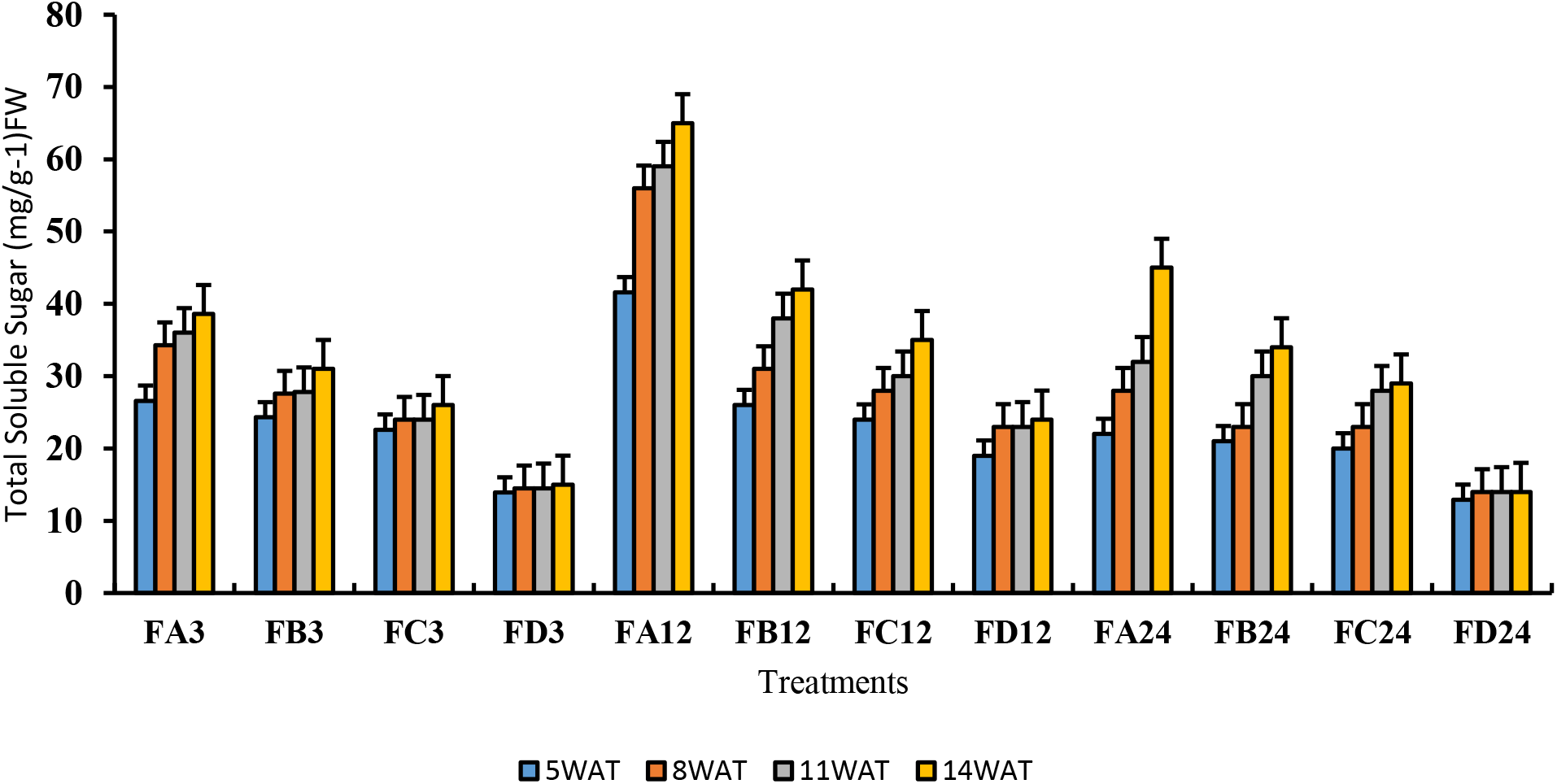
Total Soluble Sugar levels of rice plant from 3days after priming to 14 weeks after priming.

**Fig. 10:**
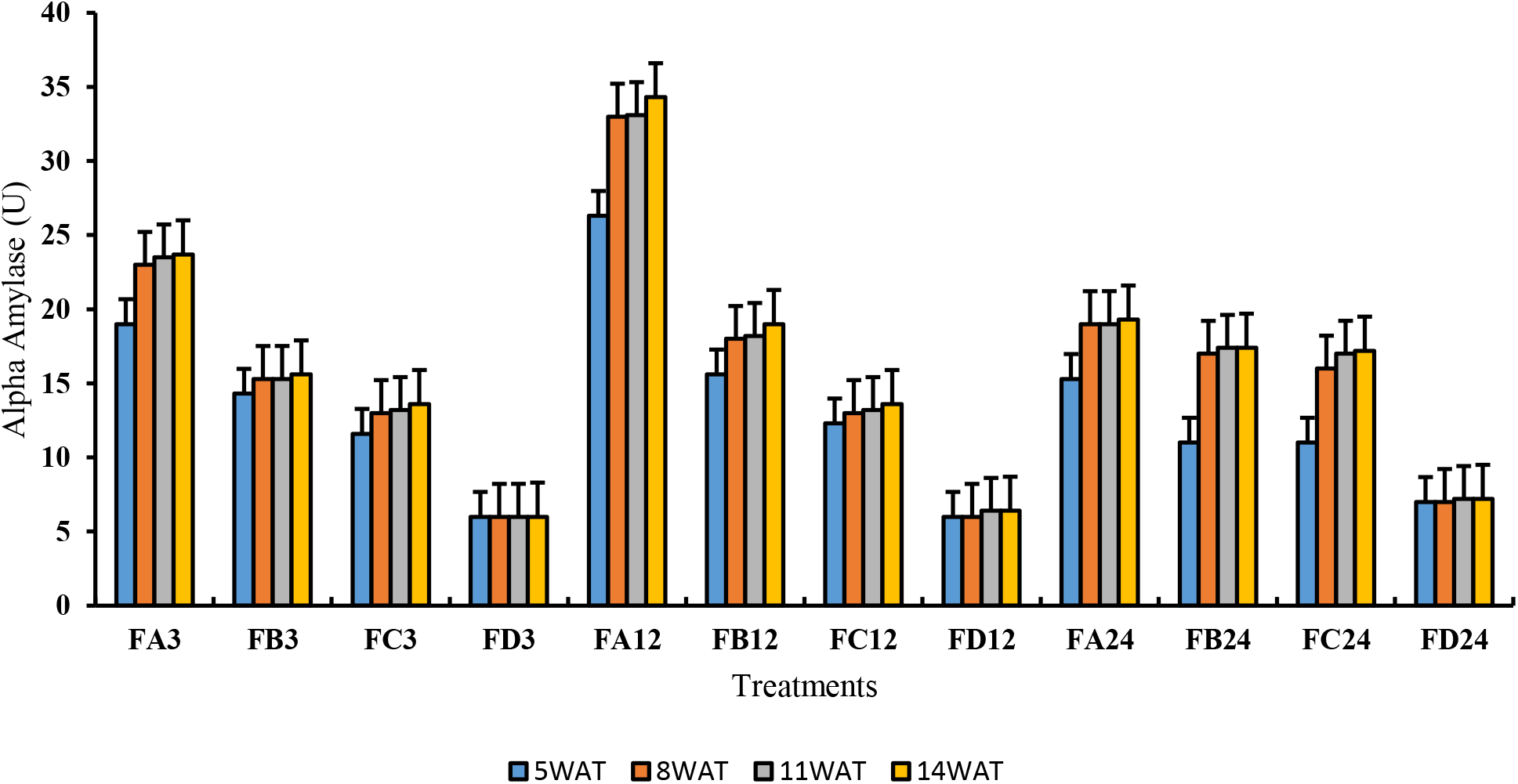
Alpha Amylase levels of rice plant from 3days after priming to 14 weeks after priming.

**Fig. 11:**
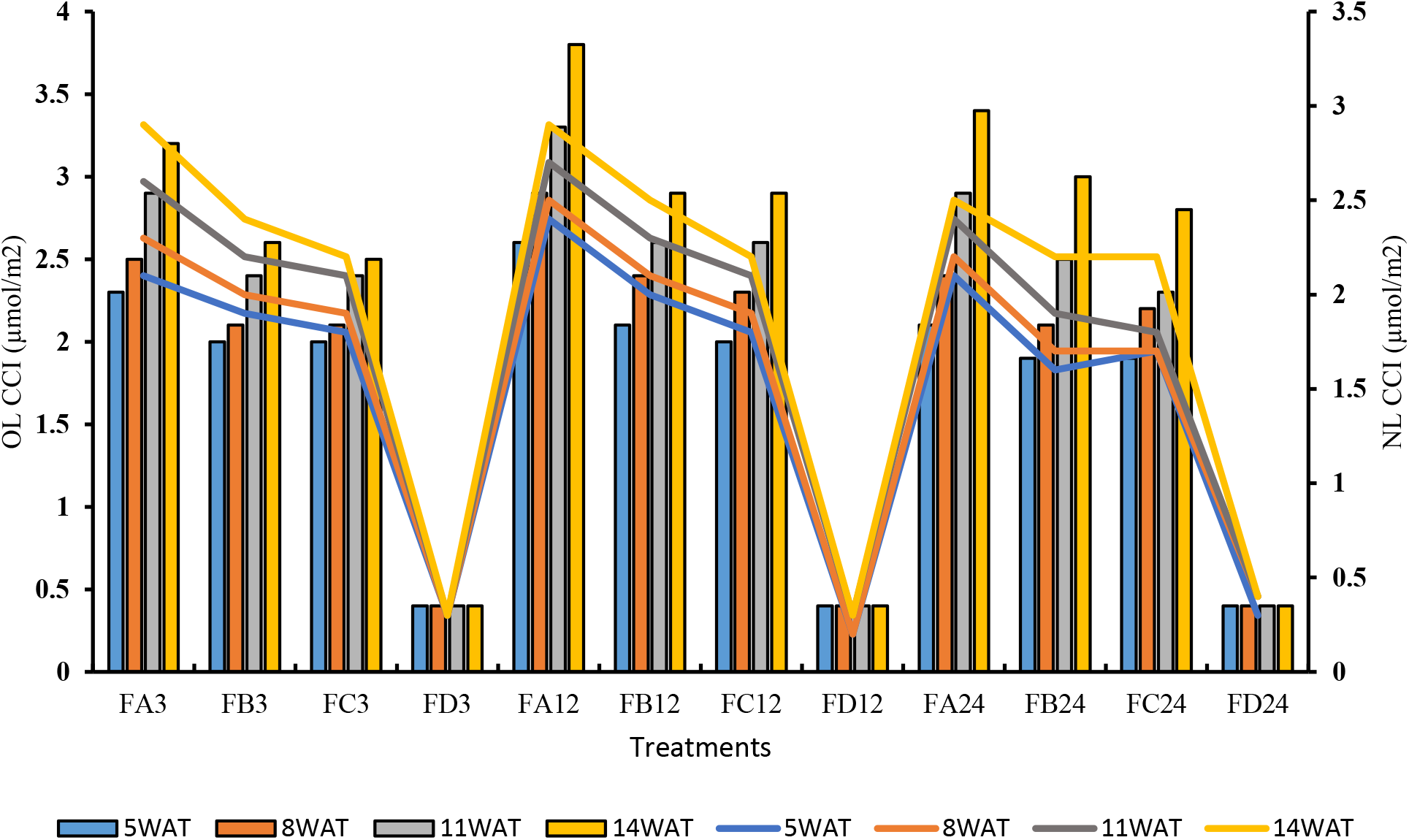
Old and New Leaf Chlorophyll Content Index of rice plant from 3days after priming to 14 weeks after priming.

The chlorophyll content of rice leaves from 5WAT to 164WAT as observed in (Table 4) showed that there were significant changes in the chlorophyll pattern. In chlorophyll-a, the non-primed rice plant was observed to be the lowest as compared to the one from the bio-primed set up. A similar observation was seen in chlorophyll-b. Generally, the BPS was observed to show highest levels of chlorophyll contents.

**Table 4.** Chlorophyll-a and Chlorophyll-b content of rice plant after bio-priming

| Samples | Chlorophyll-a (mg/cm<sup>2</sup>) FW on |  |  |  | Chlorophyll-b (mg/cm<sup>2</sup>) FW on |  |  |  |
| --- | --- | --- | --- | --- | --- | --- | --- | --- |
|  | 5WAT | 8WAT | 11WAT | 14WAT | 5WAT | 8WAT | 11WAT | 14WAT |
| FA3 | 33.7<sup>a</sup> | 38.4<sup>a</sup> | 43.3<sup>a</sup> | 49.1<sup>a</sup> | 16.2<sup>a</sup> | 16.7<sup>a</sup> | 17.0<sup>a</sup> | 17.6<sup>a</sup> |
| FB3 | 31.0<sup>b</sup> | 33.9<sup>b</sup> | 35.6<sup>b</sup> | 38.5<sup>b</sup> | 12.9<sup>b</sup> | 13.2<sup>b</sup> | 13.3<sup>b</sup> | 13.5<sup>b</sup> |
| FC3 | 31.3<sup>b</sup> | 32.8<sup>c</sup> | 34.4<sup>b</sup> | 36.8<sup>c</sup> | 12.4<sup>b</sup> | 12.6<sup>c</sup> | 12.7<sup>c</sup> | 12.9<sup>b</sup> |
| FD3 | 6.5<sup>c</sup> | 6.9<sup>d</sup> | 7.1<sup>c</sup> | 7.7<sup>d</sup> | 4.6<sup>c</sup> | 4.7<sup>d</sup> | 4.7<sup>d</sup> | 4.8<sup>c</sup> |
| FA12 | 37.4<sup>a</sup> | 45.1<sup>a</sup> | 48.2<sup>a</sup> | 53.2<sup>a</sup> | 17.8<sup>a</sup> | 18.3<sup>a</sup> | 19.0<sup>a</sup> | 19.8<sup>a</sup> |
| FB12 | 32.5<sup>b</sup> | 35.0<sup>b</sup> | 37.1<sup>b</sup> | 42.1<sup>b</sup> | 12.9<sup>b</sup> | 13.1<sup>b</sup> | 14.3<sup>b</sup> | 14.6<sup>b</sup> |
| FC12 | 31.8<sup>b</sup> | 34.2<sup>b</sup> | 36.4<sup>b</sup> | 39.2<sup>c</sup> | 12.9<sup>b</sup> | 13.0<sup>b</sup> | 13.8<sup>b</sup> | 14.0<sup>b</sup> |
| FD12 | 6.4<sup>c</sup> | 6.6<sup>c</sup> | 6.9<sup>c</sup> | 7.2<sup>d</sup> | 4.5<sup>c</sup> | 4.5<sup>c</sup> | 4.5<sup>c</sup> | 4.5<sup>c</sup> |
| FA24 | 35.1<sup>a</sup> | 39.4<sup>a</sup> | 42.1<sup>a</sup> | 48.2<sup>a</sup> | 15.6<sup>a</sup> | 15.9<sup>a</sup> | 16.5<sup>a</sup> | 16.8<sup>a</sup> |
| FB24 | 33.1<sup>b</sup> | 36.2<sup>b</sup> | 36.2<sup>b</sup> | 39.2<sup>b</sup> | 12.6<sup>b</sup> | 12.7<sup>b</sup> | 13.1<sup>b</sup> | 13.2<sup>b</sup> |
| FC24 | 33.0<sup>b</sup> | 35.1<sup>b</sup> | 35.4<sup>b</sup> | 37.1<sup>c</sup> | 12.4<sup>b</sup> | 12.5<sup>b</sup> | 13.0<sup>b</sup> | 13.1<sup>b</sup> |
| FD24 | 6.6<sup>c</sup> | 6.7<sup>c</sup> | 7.0<sup>c</sup> | 7.3<sup>d</sup> | 4.4<sup>c</sup> | 4.4<sup>c</sup> | 4.5<sup>c</sup> | 4.6<sup>c</sup> |

Results with similar alphabetic superscripts on same column did not differ from each other ( $p>0.05$ ). Results were in mean and standard error of five replicates. DAT=Day after transplant, FA3 = Ferruginous soil inoculated with *Bacillus*
sp at 3hours of bio-priming, FB3 = Ferruginous soil inoculated with *Proteus* sp at 3hours of bio-priming, FC3 = Ferruginous soil inoculated with *Klebsiella* sp. at 3hours of bio-priming. FD= Ferruginous soil without inoculation at 3hours of bio-priming. The number after each sample signifies time of priming.

### Effect of bio-primed PSB on rice biomass parameters

The biomass performance of rice plant was observed to show significant increase in the bio-primed set up compared to the non-primed set up. The BPS was observed to have higher weight of fresh plant and leaf area (Table 5) at all durations across all assayed days as against the PPS and KPS. The 12 hours priming duration for *Bacillus cereus* strain GGBSU-1 was observed to be more effective with highest biomass properties. There was no significant difference between the leaf area observed in PPS and KPS. Increased leaf number was observed in the bio-primed set up as against the non-primed seeds (Table 6). Greater percentage of the leaves observed in the non-primed set up showed signs of chlorosis and necrosis at the leaf tip (80%), while only (12%) leaf showed signs of leaf tip necrosis in the BPS at 12 hours duration (Table 6).

**Table 5.** Weight of fresh plant and area of rice leaf after bio-priming

| Samples | Weight of Fresh Plant (g<sup>-1</sup>) on |  |  |  | Largest Leaf Area (cm<sup>2</sup>) on |  |  |  |
| --- | --- | --- | --- | --- | --- | --- | --- | --- |
|  | 5WAT | 8WAT | 11WAT | 14WAT | 5WAT | 8WAT | 11WAT | 14WAT |
| FA3 | 3.5<sup>a</sup> | 3.8<sup>a</sup> | 4.2<sup>a</sup> | 4.5<sup>a</sup> | 3.2<sup>a</sup> | 3.5<sup>a</sup> | 4.0<sup>a</sup> | 4.2<sup>a</sup> |
| FB3 | 3.0<sup>b</sup> | 3.2<sup>b</sup> | 3.3<sup>b</sup> | 3.7<sup>b</sup> | 2.6<sup>b</sup> | 2.8<sup>b</sup> | 3.0<sup>b</sup> | 4.0<sup>b</sup> |
| FC3 | 3.0<sup>b</sup> | 3.1<sup>b</sup> | 3.2<sup>b</sup> | 3.5<sup>c</sup> | 2.5<sup>b</sup> | 2.9<sup>b</sup> | 3.1<sup>b</sup> | 3.8<sup>b</sup> |
| FD3 | 1.0<sup>c</sup> | 1.1<sup>c</sup> | 1.2<sup>c</sup> | 1.3<sup>d</sup> | 0.9<sup>c</sup> | 1.0<sup>c</sup> | 1.1<sup>c</sup> | 1.2<sup>c</sup> |
| FA12 | 3.9<sup>a</sup> | 4.3<sup>a</sup> | 4.8<sup>a</sup> | 5.2<sup>a</sup> | 3.7<sup>a</sup> | 4.0<sup>a</sup> | 4.3<sup>a</sup> | 4.8<sup>a</sup> |
| FB12 | 3.7<sup>b</sup> | 3.9<sup>b</sup> | 4.2<sup>b</sup> | 4.6<sup>b</sup> | 3.6<sup>b</sup> | 3.8<sup>b</sup> | 4.0<sup>b</sup> | 4.3<sup>b</sup> |
| FC12 | 3.6<sup>b</sup> | 3.8<sup>b</sup> | 3.9<sup>b</sup> | 4.3<sup>c</sup> | 3.4<sup>b</sup> | 3.6<sup>b</sup> | 3.7<sup>c</sup> | 4.2<sup>b</sup> |
| FD12 | 1.0<sup>c</sup> | 1.2<sup>c</sup> | 1.3<sup>c</sup> | 1.4<sup>d</sup> | 1.0<sup>c</sup> | 1.0<sup>c</sup> | 1.1<sup>d</sup> | 1.2<sup>c</sup> |
| FA24 | 3.8<sup>a</sup> | 4.0<sup>a</sup> | 4.4<sup>a</sup> | 4.6<sup>a</sup> | 3.6<sup>a</sup> | 3.8<sup>a</sup> | 4.2<sup>a</sup> | 4.4<sup>a</sup> |
| FB24 | 3.7<sup>a</sup> | 3.9<sup>a</sup> | 4.2<sup>b</sup> | 4.5<sup>a</sup> | 3.6<sup>a</sup> | 3.7<sup>a</sup> | 4.0<sup>a</sup> | 4.4<sup>a</sup> |
| FC24 | 3.6<sup>a</sup> | 3.8<sup>a</sup> | 4.0<sup>c</sup> | 4.3<sup>b</sup> | 3.5<sup>a</sup> | 3.7<sup>a</sup> | 3.9<sup>a</sup> | 4.2<sup>a</sup> |
| FD24 | 1.0<sup>b</sup> | 1.1<sup>b</sup> | 1.2<sup>d</sup> | 1.3<sup>c</sup> | 1.0<sup>b</sup> | 1.0<sup>b</sup> | 1.1<sup>b</sup> | 1.2<sup>b</sup> |

Results with similar alphabetic superscripts on same column did not differ from each other (p>0.05). Results were in mean and standard error of five replicates. WAT=Weeks after transplant, FA3 = Ferruginous soil inoculated with *Bacillus* sp at 3hours of bio-priming, FB3 = Ferruginous soil inoculated with *Proteus* sp at 3hours of bio-priming, FC3 = Ferruginous soil inoculated with *Klebsiella* sp. at 3hours of bio-priming. FD= Ferruginous soil without inoculation at 3hours of bio-priming. The number after each sample signifies time of priming.

**Table 6.**
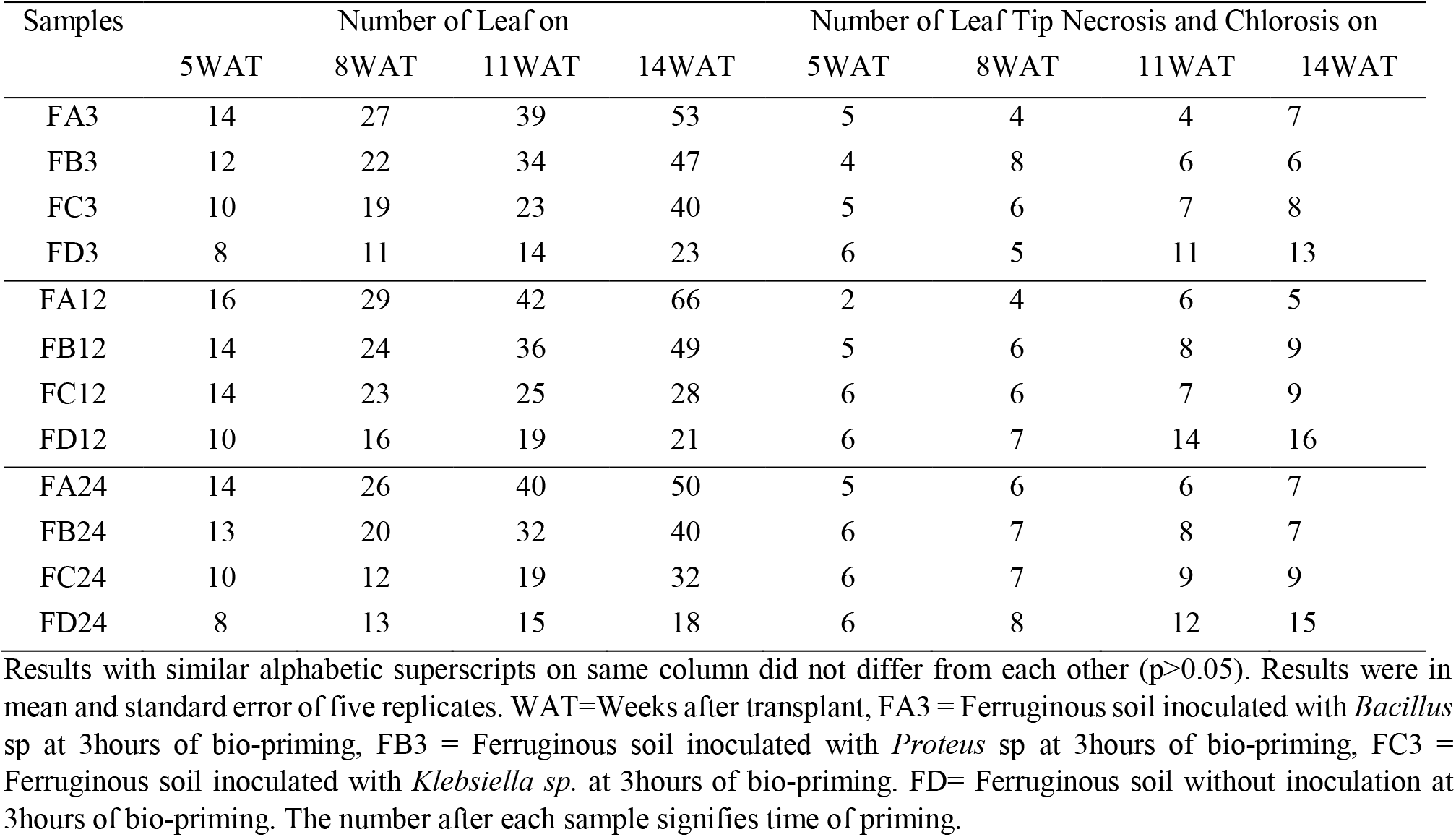
Number of rice leaf and number of leaf with signs of necrosis and chlorosis at the tip after bio-priming

### Effect of bio-primed PSB on rice root parameters

Fresh root length and dry root length (Table 7; Table 8) also revealed improved parameter in the FA12 than all other bio-prime set up. The number of secondary root (Table 7) was observed to increase in all the assayed pots. High number of secondary root was observed in the ferruginous soil without inoculation at 3 weeks after transplanting (3WAT) at all bio-priming duration. However, this was reduced significantly with 40% at 6WAT and 8WAT.

**Table 7.**
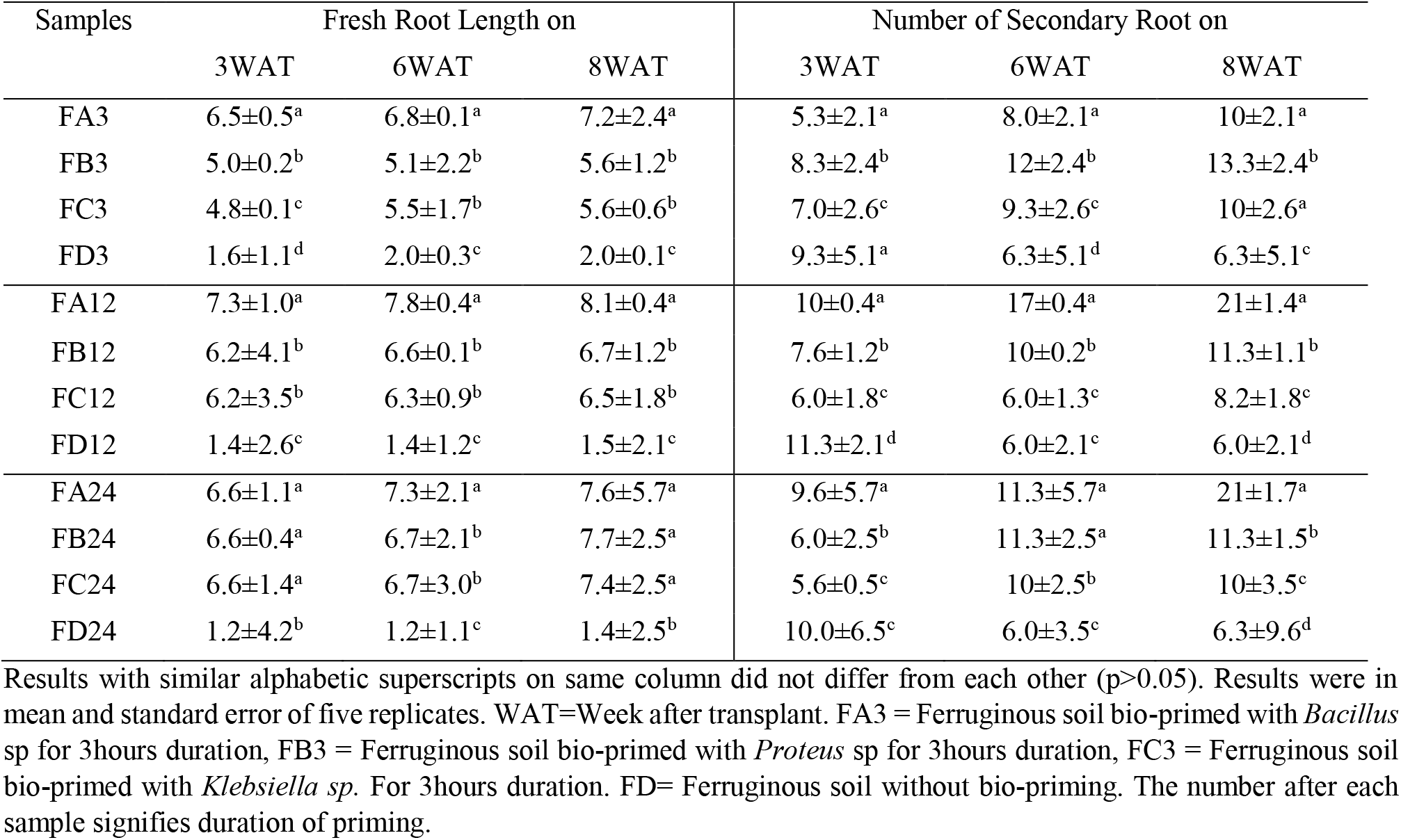
Response of fresh root length and number of secondary roots of rice plant to PSB bio-priming.

| Samples | Fresh Root Length on |  |  | Number of Secondary Root on |  |  |
| --- | --- | --- | --- | --- | --- | --- |
|  | 3WAT | 6WAT | 8WAT | 3WAT | 6WAT | 8WAT |
| FA3 | 6.5±0.5<sup>a</sup> | 6.8±0.1<sup>a</sup> | 7.2±2.4<sup>a</sup> | 5.3±2.1<sup>a</sup> | 8.0±2.1<sup>a</sup> | 10±2.1<sup>a</sup> |
| FB3 | 5.0±0.2<sup>b</sup> | 5.1±2.2<sup>b</sup> | 5.6±1.2<sup>b</sup> | 8.3±2.4<sup>b</sup> | 12±2.4<sup>b</sup> | 13.3±2.4<sup>b</sup> |
| FC3 | 4.8±0.1<sup>c</sup> | 5.5±1.7<sup>b</sup> | 5.6±0.6<sup>b</sup> | 7.0±2.6<sup>c</sup> | 9.3±2.6<sup>c</sup> | 10±2.6<sup>a</sup> |
| FD3 | 1.6±1.1<sup>d</sup> | 2.0±0.3<sup>c</sup> | 2.0±0.1<sup>c</sup> | 9.3±5.1<sup>a</sup> | 6.3±5.1<sup>d</sup> | 6.3±5.1<sup>c</sup> |
| FA12 | 7.3±1.0<sup>a</sup> | 7.8±0.4<sup>a</sup> | 8.1±0.4<sup>a</sup> | 10±0.4<sup>a</sup> | 17±0.4<sup>a</sup> | 21±1.4<sup>a</sup> |
| FB12 | 6.2±4.1<sup>b</sup> | 6.6±0.1<sup>b</sup> | 6.7±1.2<sup>b</sup> | 7.6±1.2<sup>b</sup> | 10±0.2<sup>b</sup> | 11.3±1.1<sup>b</sup> |
| FC12 | 6.2±3.5<sup>b</sup> | 6.3±0.9<sup>b</sup> | 6.5±1.8<sup>b</sup> | 6.0±1.8<sup>c</sup> | 6.0±1.3<sup>c</sup> | 8.2±1.8<sup>c</sup> |
| FD12 | 1.4±2.6<sup>c</sup> | 1.4±1.2<sup>c</sup> | 1.5±2.1<sup>c</sup> | 11.3±2.1<sup>d</sup> | 6.0±2.1<sup>c</sup> | 6.0±2.1<sup>d</sup> |
| FA24 | 6.6±1.1<sup>a</sup> | 7.3±2.1<sup>a</sup> | 7.6±5.7<sup>a</sup> | 9.6±5.7<sup>a</sup> | 11.3±5.7<sup>a</sup> | 21±1.7<sup>a</sup> |
| FB24 | 6.6±0.4<sup>a</sup> | 6.7±2.1<sup>b</sup> | 7.7±2.5<sup>a</sup> | 6.0±2.5<sup>b</sup> | 11.3±2.5<sup>a</sup> | 11.3±1.5<sup>b</sup> |
| FC24 | 6.6±1.4<sup>a</sup> | 6.7±3.0<sup>b</sup> | 7.4±2.5<sup>a</sup> | 5.6±0.5<sup>c</sup> | 10±2.5<sup>b</sup> | 10±3.5<sup>c</sup> |
| FD24 | 1.2±4.2<sup>b</sup> | 1.2±1.1<sup>c</sup> | 1.4±2.5<sup>b</sup> | 10.0±6.5<sup>c</sup> | 6.0±3.5<sup>c</sup> | 6.3±9.6<sup>d</sup> |

Results with similar alphabetic superscripts on same column did not differ from each other ( $p>0.05$ ). Results were in mean and standard error of five replicates. WAT=Week after transplant. FA3 = Ferruginous soil bio-primed with *Bacillus* sp for 3hours duration, FB3 = Ferruginous soil bio-primed with *Proteus* sp for 3hours duration, FC3 = Ferruginous soil bio-primed with *Klebsiella* sp. For 3hours duration. FD= Ferruginous soil without bio-priming. The number after each sample signifies duration of priming.

**Table 8.** Response of dry root length of rice plant to PSB bio-priming.

| Samples | Dry Root Length (cm) on |  |  |
| --- | --- | --- | --- |
|  | 3WAT | 6WAT | 8WAT |
| FA3 | 5.3±3.1<sup>a</sup> | 5.9±2.1<sup>a</sup> | 6.3±2.1<sup>a</sup> |
| FB3 | 4.1±1.4<sup>b</sup> | 4.3±2.4<sup>b</sup> | 4.7±2.4<sup>b</sup> |
| FC3 | 4.0±2.6<sup>b</sup> | 4.1±2.6<sup>b</sup> | 4.5±2.6<sup>b</sup> |
| FD3 | 0.6±5.1<sup>c</sup> | 0.6±5.1<sup>c</sup> | 0.6±5.1<sup>c</sup> |
| FA12 | 6.2±1.4<sup>a</sup> | 6.6±0.4<sup>a</sup> | 7.0±1.4<sup>a</sup> |
| FB12 | 5.5±1.2<sup>b</sup> | 5.7±0.2<sup>b</sup> | 5.9±1.1<sup>b</sup> |
| FC12 | 5.1±1.8<sup>c</sup> | 5.4±1.3<sup>b</sup> | 5.6±1.8<sup>c</sup> |
| FD12 | 0.6±2.1<sup>d</sup> | 0.9±2.1<sup>c</sup> | 1.0±2.1<sup>d</sup> |
| FA24 | 6.0±5.7<sup>a</sup> | 6.4±5.7<sup>a</sup> | 6.7±1.7<sup>a</sup> |
| FB24 | 5.3±2.5<sup>b</sup> | 5.6±2.5<sup>b</sup> | 6.0±1.5<sup>b</sup> |
| FC24 | 5.3±0.5<sup>c</sup> | 5.4±2.5<sup>b</sup> | 5.8±3.5<sup>c</sup> |
| FD24 | 0.7±6.5<sup>d</sup> | 1.0±3.5<sup>c</sup> | 1.0±9.6<sup>d</sup> |

Results with similar alphabetic superscripts on same column did not differ from each other ( $p>0.05$ ). Results were in mean and standard error of five replicates. WAT=Week after transplant. FA3 = Ferruginous soil bio-primed with *Bacillus* sp for 3hours duration, FB3 = Ferruginous soil bio-primed with *Proteus* sp for 3hours duration, FC3 = Ferruginous soil bio-primed with *Klebsiella* sp. For 3hours duration. FD= Ferruginous soil without bio-priming. The number after each sample signifies duration of priming.

### General growth characteristics

The bio-primed set up was observed to achieve maturity at 11^th^ week after bio-priming. However, for the ferruginous soil without priming with PSBs (FD), growth and yield seized at the 8^th^ week after priming (Fig. 12).

**Fig. 12:**
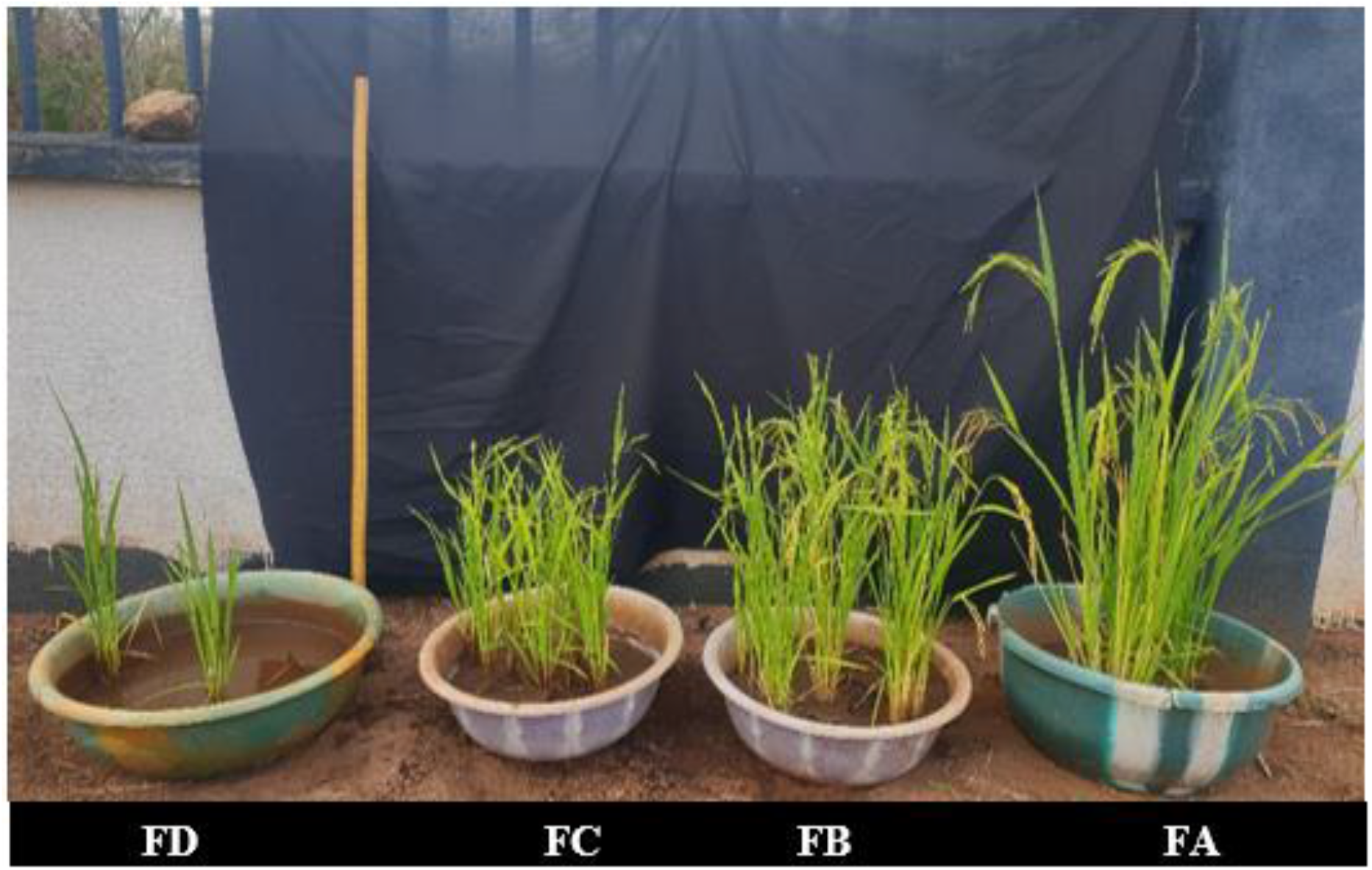
Growth performance of rice at 8^th^ weeks after bio-priming.

Fig. 13 showed the number of tillers and reproductive tillers observed at harvest day. Bio-priming of rice seeds has significantly improved the number of tillers and the number of reproductive tillers in rice plants at harvest. The BPS was observed to show highest NT and NRT at all priming duration as compared to the KPS and BPS. The 12 hours priming duration was observed to show highest growth yield properties at harvest. The ferruginous soil without priming stopped growing before the time at which tillers appeared. Similarly, higher number of panicle per plant and seed per panicle (Fig. 14) were observed in the PSB primed set up, while the rice plant in ferruginous soil without bio-priming did not survive till this stage.

**Fig. 13:**
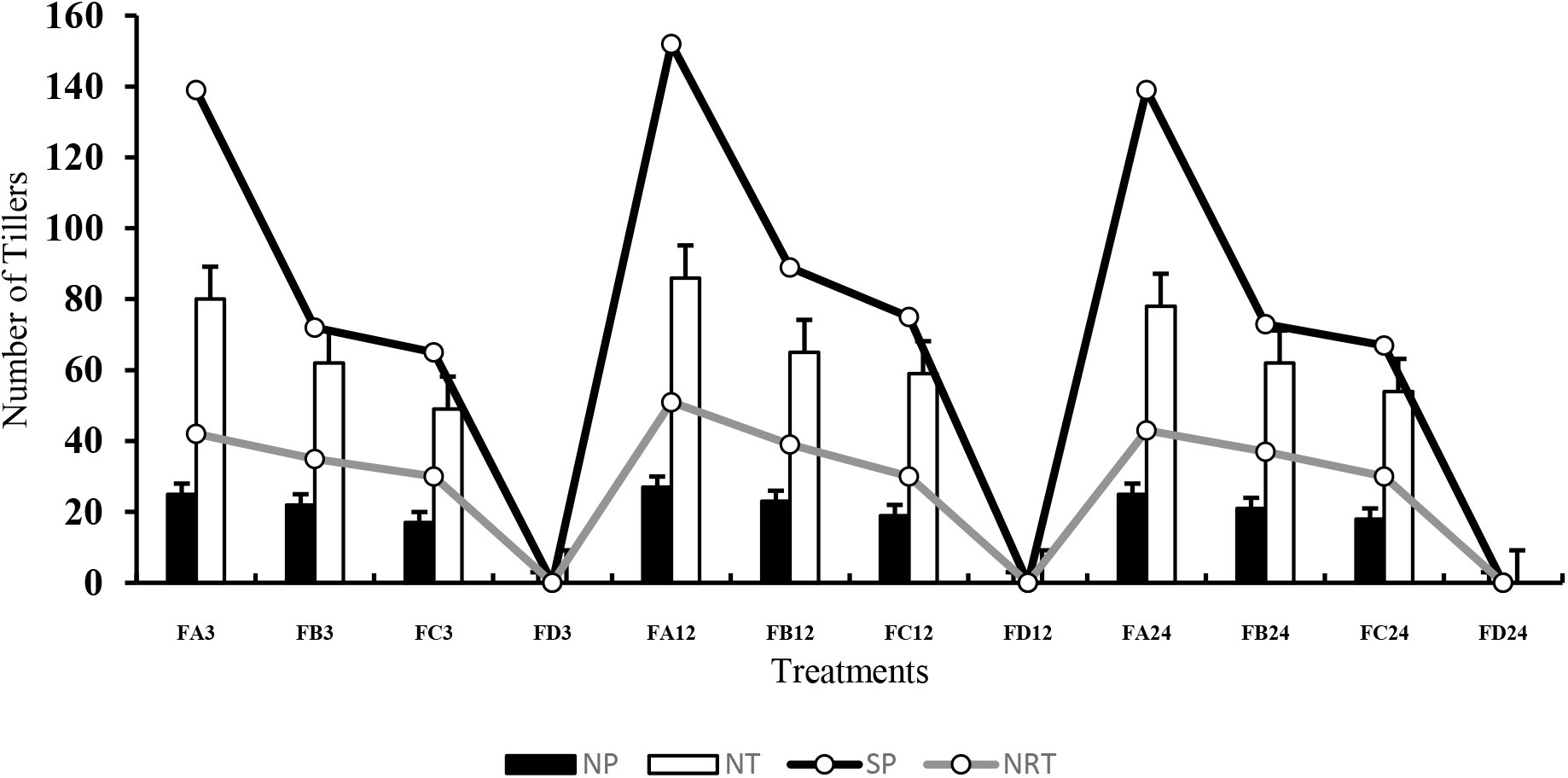
FA-FC= Ferruginous soil under bio-priming experiment. With A, B and C bacteria respectively, A= *Bacillus cereus strain GGBSU-1*. B = *Proteus mirabilis* strain TL14-1. respectively. C = *Klebsiella variicola* strain AUH-KAM-9.

**Fig. 14:**
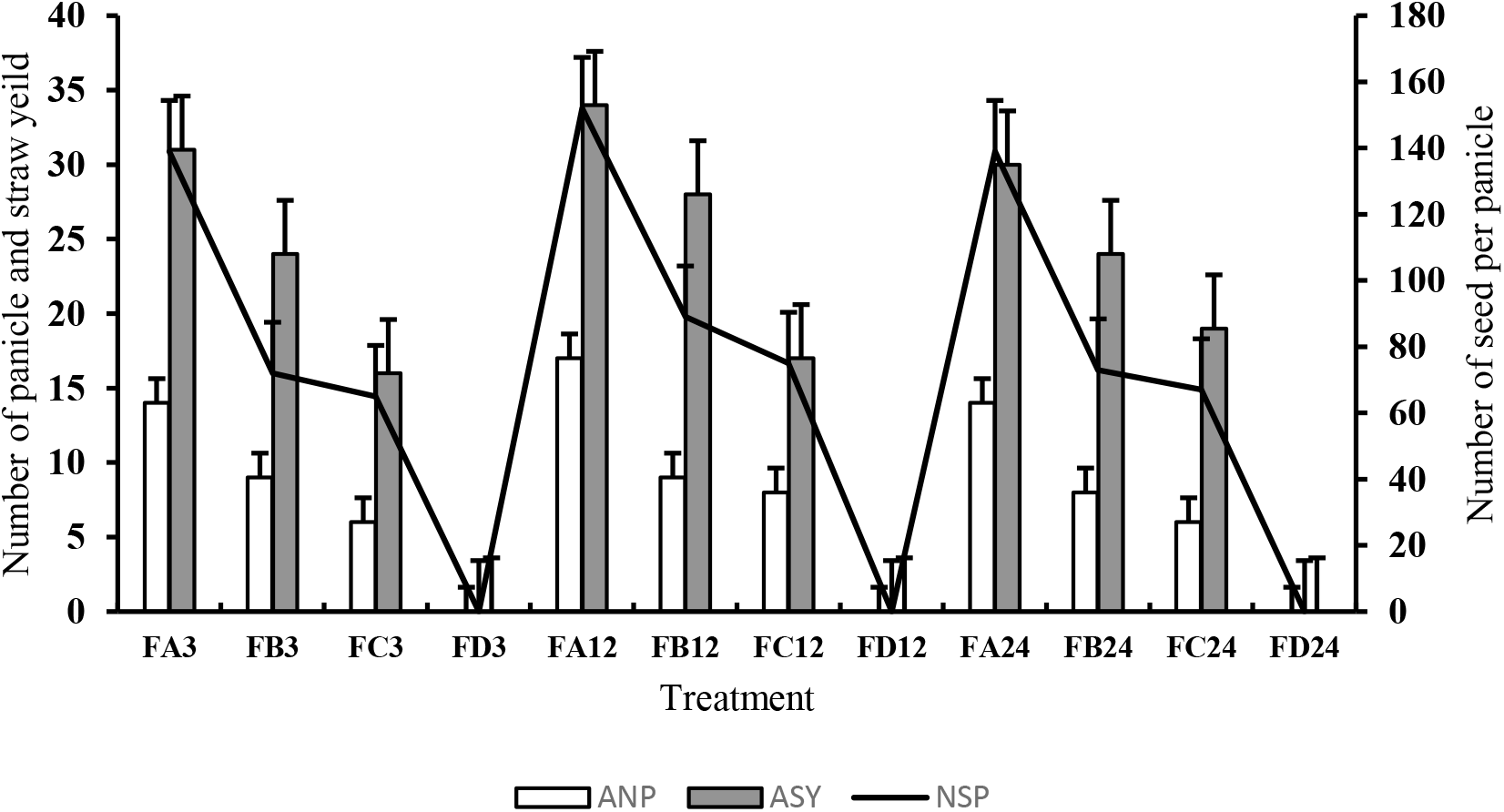
ANP =Number of panicle, ASY =Straw yield and NSP =Number of seed per panicle. FA-FC= Ferruginous soil under bio-priming experiment With A, B and C bacteria respectively, A= *Bacillus cereus strain GGBSU-1*. B = *Proteus mirabilis* strain TL14-1 respectively. C = *Klebsiella variicola* strain AUH-KAM-9.

Furthermore, rice yield parameters at harvest (Table 9) revealed that weight of rice panicle is higher in the BPS (FA), especially at 12 hours of bio-priming as against the other priming durations. The 100-grain weight at harvest was observed to show no significant difference between the KPS (FB) and PPS (FC). About 5% increase in the 100-grain weight was observed in the BIS (FA). A similar results was obtained in the plant tissue water content (Table 19). The BIS (FA) at 12 hours duration was observed to have highest yield in terms of weight of peduncle without rice, weight of husked and de-husked rice seed (Table 9 and Fig. 15).

**Table 9.** Rice yield parameters at harvest

| Soil samples | WRP (g) | WPWR (g) | 100 grains (g) | PTW |
| --- | --- | --- | --- | --- |
| FA3 | 2.11 ± 1.05<sup>a</sup> | 0.59 ± 0.21<sup>a</sup> | 2.6 ± 0.07<sup>a</sup> | 0.73 ± 0.41<sup>a</sup> |
| FB3 | 2.08 ± 0.05<sup>a</sup> | 0.38 ± 0.17<sup>b</sup> | 2.5 ± 0.95<sup>a</sup> | 0.64 ± 0.24<sup>b</sup> |
| FC3 | 1.94 ± 1.10<sup>b</sup> | 0.17 ± 0.14<sup>c</sup> | 2.1 ± 0.06<sup>b</sup> | 0.64 ± 0.18<sup>b</sup> |
| FA12 | 2.69 ± 0.05<sup>a</sup> | 0.87 ± 0.11<sup>a</sup> | 2.7 ± 0.07<sup>a</sup> | 0.78 ± 0.11<sup>a</sup> |
| FB12 | 2.58 ± 0.02<sup>b</sup> | 0.70 ± 0.89<sup>b</sup> | 2.6 ± 0.05<sup>a</sup> | 0.76 ± 0.21<sup>a</sup> |
| FC12 | 2.54 ± 0.10<sup>b</sup> | 0.23 ± 0.04<sup>c</sup> | 2.4 ± 0.06<sup>a</sup> | 0.69 ± 0.23<sup>b</sup> |
| FA24 | 2.08 ± 0.07<sup>a</sup> | 0.58 ± 0.11<sup>a</sup> | 2.6 ± 0.11<sup>a</sup> | 0.72 ± 0.81<sup>a</sup> |
| FB24 | 2.06 ± 0.87<sup>a</sup> | 0.38 ± 0.07<sup>b</sup> | 2.6 ± 0.50<sup>a</sup> | 0.64 ± 0.14<sup>b</sup> |
| FC24 | 1.90 ± 0.05<sup>b</sup> | 0.12 ± 0.08<sup>c</sup> | 2.1 ± 0.12<sup>c</sup> | 0.65 ± 0.08<sup>b</sup> |

WRP= Weight of rice panicle, WPWR = weight of peduncle without rice, 100 grains = weight of 100 grains and PTW= plant tissue water. Results with similar alphabetic superscripts on same column did not differ from each other (p>0.05). FA-FD= Ferruginous soil, NF= Non-ferruginous soil.

**Fig. 15:**
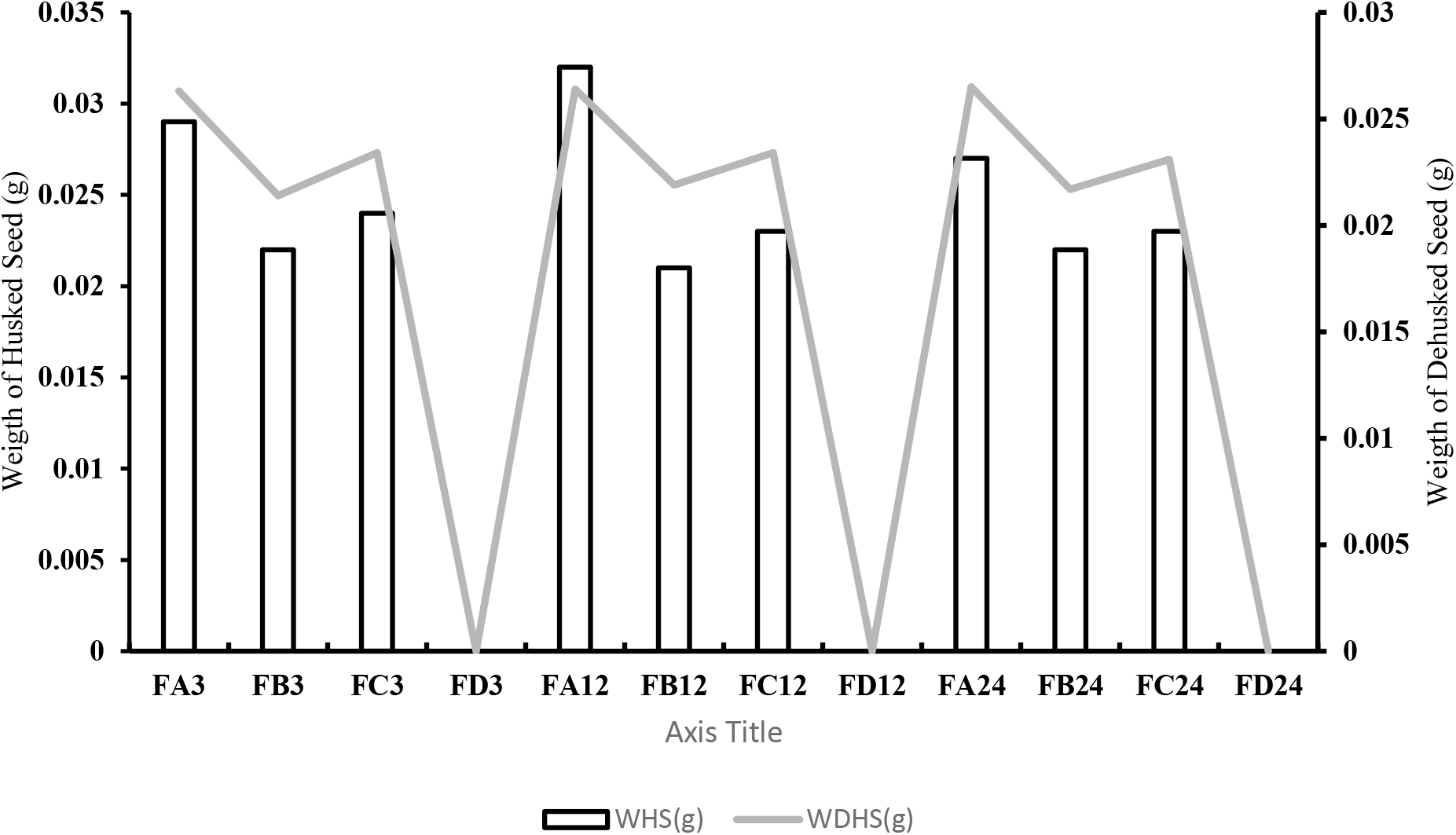
Weight of husked seed (WHS) and weight of dehusked seed (WDHS) at harvest day. Results are average of five replicates. FA-FC= Ferruginous soil inoculated with A, B and C bacteria respectively, NF= Non-ferruginous soil. A= *Bacillus cereus strain GGBSU-1*. B = *Proteus mirabilis* strain TL14-1 respectively. C = *Klebsiella variicola* strain AUH-KAM-9

## Discussion

Possibilities to improve the growth parameters of rice plant through bio-priming with phosphate solubilizing bacteria under ferruginous ultisol condition was analyzed. Ferruginous ultisol is a special soil condition with high acidity, high iron levels, low bioavailable phosphorus and low water holding capacity (Musa and Ikhajiagbe, 2020a). The physico-chemical results of the soil (Table 1) used in this study showed similar parameters as the condition described above, which is regarded by Musa and Ikhajigbe (2021) and Wang *et al*. (2014) as a soil with poor growing condition because of its ferruginicity nature. Since the current study relied on bio-priming of a phosphate solubilizing bacteria with rice seed, biosafety analysis of the bacteria indicated that the bacteria are not pathogenic to humans and therefore safe for further plant yield studies. This is related with the work of Muhamad et al. (2020), who confirmed the hemolysis of certain phosphorus and potassium solubilizing *Enterobacter* sp as inoculant in crop improvement.

The fast coleoptile appearance observed in the rice seeds that were bio-primed with the PSBs proved the effectiveness of bio-priming with PSB in improving germination properties as against the non-primed seeds. This is likely by stimulating certain metabolites such as α-amylase which improves coleoptile growth (Setter and Waters, 2003), thus increase chances of survival for the growing seedling (Huang et al., 2005). Furthermore, the fast coleoptile appearance resulted in early germination in the bio-primed seeds at 2 days after transplanting as against the seeds not bio-primed. This finding is consistent with an *in vitro* plate bioassay conducted by Suleman et al., (2018) when they reported that *Enterobacter spp*. (MS32P) increased the germination properties of wheat. Similarly, seed bio-priming with strains of endophytic bacteria on muskmelon seeds resulted in the promotion of hypocotyl and radicle growth (Duan et al., 2013). Also, the 12 hours seed priming with the bacterial isolates proved more effective than the 3 and 24 hours. This is consistent with the study by Ananthi et al. (2014) who investigated the effects of different durations of bio-priming chilli seeds with *Trichoderma viride* and discovered that bio-priming for 12 hours is the most effective duration with improved germination percentage and germination rate. This may be that at 24 hours, the bacterial population in seed became overwhelming and therefore slow down the energy efficiency of the seed (Basavaraj et al., 2019). According to Anitha et al., 2013, Germination and enhanced seedling establishments can be obtained through seed bio-priming with plant growth promoting (PGP) bacteria.

Since bio-priming with PSB bacteria having PGP capabilities can change the seed microbiome and bring about root colonization in soils (Gupta *et al*., 2012a), effective bacteria colonization in rice seed endosperm was observed using SEM where a large number of bacterial cells were detected on the rice seed endosperm. This showed that these PSBs are motile and can easily proliferate within the rhizospheric environment and possibly influence the Fe-P bond (Sachdev *et al*., 2009).

As the rice seedling grows in the ferruginous soil, the improved morphological parameters such as shoot length and length of internodes observed in the bio-primed set up as against the non-primed set up showed the potential of the PSBs in improving growth conditions of rice plant, even under ferruginous condition with the FA12 hours priming duration showing highest influence even though the soils have poor growing conditions (Table 1) and has been documented by Ikhajiagbe et al., 2021 to bring about metabolic disorders in plants, leading to reduced plant yield as observed in the seeds without bio-priming. According to Sharifi, (2011), a significance increase in crop growth, dry matter accumulation and grain yield were observed in maize when bio-primed with different strains of *Azotobacter* and *Azospirillum spp. Trichoderma harzianum*, *T. viride*, *B. subtilus* and *P. flouresens* as bio-priming agents of soybean seeds. A significant increase in plant height, shoot weight, number of branches and resistance against root rot disease as well as seedling protection against soil borne infection were observed (Mona et al., 2017). Since Suleman et al. (2018) highlighted a direct connection between the bacterial pathway for P-solubilization and the secretion of multiple cellular organic acids, it may be that the low molecular weight organic matter produced by the PSBs have lowered soil pH and chelated the Fe-P bound and thereby improving P availability, which in turns improve soil properties (Musa and Ikhajiagbe 2020; Zhao, 2014). The high number of secondary roots observed in the non-primed set up even though with the least root length may indicate nutrient deficiencies. Plants are known to show different responses to different specific nutrient deficiencies and the responses can vary between species. An example of such response is the secretion of a metabolite such as terpenoid (Musa and ikhajiagbe, 2021; Esha et al., 2017). This may be the reason why higher terpanoid was observed in the non-primed seeds. According to Jones and Ljung (2012), the most common changes in plants due to nutrient deficiencies are inhibition of primary root growth (often associated with P deficiency) and increase in secondary roots density (often associated with P deficiency and Fe toxicity).

The higher CCI observed in the bio-primed setup showed the positive influence of PSBs as against the un-primed set up. Since nutrient availability is considered a limiting step in plant growth and nutrition (Rodriguez and Fraga, 2009), this evidence suggests a significant contribution of the PSB to promote the rice plant photosynthetic properties and yield. Kennedy et al. (2004) observed that wide range of PGP bacteria can influence the photosynthetic signals of plants, resulting in improved grain yield and vegetative growth. Additionally, a recent review of bio-priming techniques validated that the major microbial contributions towards enhanced plant productivity included and not limited to, N2 fixation, P-soubilization, secretion of plant growth promoting hormones, secretion of beneficial organic acids, enhancement of nutrient uptake by the rhizosphere and iron chelation (Ikhajiagbe and Musa, 2020b; Mahmood and Kataoka, 2018) where all these have been observed in the present study. The 12 hours bio-priming was observed to show highest physiological properties, this indicated the influence of priming duration on rice growth. Priming duration and water potential of priming agents are considered as factors influencing priming (Ashraf and Foolad, 2005; Abdulrahmani et al., 2007).

Evidence from the rice plant biomass parameters suggests a significant contribution of the PSB in improving weight of rice plant, leaf area, number of leaf and leaf health as against the non-primed rice plant. This may be because of the growth promoting properties of the PSBs and also their ability to solubilize P in soil and make it bioavailable, since P is considered as a limiting nutrient for rice plant (Doberman and Fairhurst, 2000). This study is consistent with the work of Gholami et al. (2009) where they observed that stem, leaf area and total seedling of maize plant was increased with the bio-priming of PGPR in maize seedlings in a laboratory experiment. The high leaf tip chlorosis and necrosis that was observed in the non-primed setup indicated the poor growing condition of ferruginous soil due to the toxic effect of iron. Ferruginous soil with high Fe and pH have been associated with chlorosis of emerging leaves, low photosynthetic properties and stunted growth (Mahender et al., 2019) as observed in the ferruginous soil without priming in the current study. Priming of rice seed with these bacteria have significantly showed positive influence.

The stunted growth observed in the non-primed ferruginous soil may be as a results of the limited bioavailable phosphorus and iron toxicity of the ferruginous soils. Phosphorus is needed in large quantities in plants and it helps in early stages of cell division therefore, phosphorus limitation can cause death of plants or delay maturity (Uchida, 2000). The enhancement in numbers of tillers, reproductive tillers, 100 grain weight, weight of the peduncle without rice and plant tissue water by the PSBs was made possible because of the ability of the PSBs to act through several mechanisms (Singh and Singh, 2011). The root architecture in the FA showed more signs of bacterial colonization as compared to FB and FC (Figure 13), this may be that the *Bacillus cereus strain GGBSU-1* primed in the FA is more effective in colonizing rice plant rhizosphere and improving the poor growing condition of ferruginous ultisol. Since bio-priming of rice seeds with *Bacillus cereus strain GGBSU-1* at 12 hours priming duration (FA12) showed highest rice growth and yield effect under ferruginous condition, this research recommend bio-priming of other important crops such as maize with PSBs in ferruginous soil to improve productivity of grains in regions with iron toxic soil conditions.

## Conclusions

To the best of our knowledge, this is the first report to test the possibilities of improving growth and yield parameters of rice plant through bio-priming with phosphate solubilizing bacteria under high iron soil with limited bio-available phosphorus. Generally, the increase bacterial cell in the endosperm corresponded with in the improved morphological, physiological and biomass properties observed in the rice plants that were bio-primed with the PSBs under ferruginous soil conditions. Therefore, bio-priming of rice seeds with the phosphate solubilizing bacteria might be considered as a promising strategy for plant growth promotion under edaphic stress, especially the priming with *Bacillus cereus strain GGBSU-1* under 12 hours priming duration. Further studies on genomic and metabolomic facets of these species in relation to their growth-promoting capabilities and phosphate solubilization efficiencies might be important in improving strategies for sustainable crop production under different edaphic conditions.

## List of Abbreviations

FU: Ferruginous ultisol
CFU/mL: Colony forming unit per milliliter
FA3: Rice growing in ferruginous soil, Which was bio-primed with *Bacillus cereus* strain GGBSU-1 for 3 hours bio-priming duration
FA12: Rice growing in ferruginous soil, Which was bio-primed with *Bacillus cereus* strain GGBSU-1 for 12 hours bio-priming duration
FA24: Rice growing in ferruginous soil, Which was bio-primed with *Bacillus cereus* strain GGBSU-1 for 24 hours bio-priming duration
FB3: Rice growing in ferruginous soil, Which was bio-primed with *Proteus mirabilis* strain TL14-1 for 3 hours bio-priming duration
FB12: Rice growing in ferruginous soil, Which was bio-primed with *Proteus mirabilis* strain TL14-1 for 12 hours bio-priming duration
FB24: Rice growing in ferruginous soil, Which was bio-primed with *Proteus mirabilis* strain TL14-1 for 24 hours bio-priming duration
FC3: Rice growing in ferruginous soil, Which was bio-primed with *Klebsiella variicola* strain AUH-KAM-9 for 3 hours bio-priming duration
FC12: Rice growing in ferruginous soil, Which was bio-primed with *Klebsiella variicola* strain AUH-KAM-9 for 12 hours bio-priming duration
FC24: Rice growing in ferruginous soil, Which was bio-primed with *Klebsiella variicola* strain AUH-KAM-9 for 24 hours bio-priming duration
FD3: Rice growing in ferruginous soil without bio-priming, but soaked in water for 3 hours
FD12: Rice growing in ferruginous soil without bio-priming, but soaked in water for 12 hours
FD24: Rice growing in ferruginous soil without bio-priming, but soaked in water for 24 hours
16S rRNA: 16S ribosomal RNA
TSS: Total soluble sugar contents
AA: Alpha amylase
CCI: Chlorophyll Context Index
OL: Old Leaf
NL: New Leaf
ANP: Number of Panicle per pot
ASY: Straw Yield
NSP: Number of Seed per Panicle at harvest day
WRP: Weight of Rice Panicle
WPWR: Weight of Peduncle without Rice
PTW: Plant Tissue Water
WHS: Weight of Husked Seed
WDHS: Weight of Dehusked Seed at harvest day
DAT: Days After Transplanting
WAP: Weeks After Priming
SEM: Scanning Electron Microscope
SOM: Soil Organic Matter
CEC: Cation Exchange Capacity
OC: Organic Carbon
EA: Exchangeable Acidity
PGP: Plant Growth Promotion
PSB: Phosphate Solubilizing Bacteria
T/LF: Time Taken For Leaf Formation
NL: Number of Leaves Per Unit Time
N: Number of Periods
T: Time During Which Parameter was Measured
BPS: *Bacillus cereus* strain GGBSU-1 Primed Seed
KPS: *Klebsiella variicola* strain AUH-KAM-9 Primed Seed
PPS: *Proteus mirabilis* strain TL14-1 Primed Seed.

## Acknowledgements

The researchers are grateful to the Vice-Chancellor of Admiralty University of Nigeria and its Management for their support during the field study and lab work. The support and assistance from the Department of Plant Biology and Biotechnology and the Department of Microbiology both in the University of Benin, Nigeria is well appreciated. The mentorship and efforts of my supervisor, Beckley Ikhajiagbe, Ph.D., FIPMD, of the Department of Plant Biology and Biotechnology during the course of the study is very much appreciated. My sincere thanks to the entire staff of the Department of Material Science and Engineering, Obafemi Awolowo University, Ile-ife for allowing MSI access to SEM during the study.

## Authors’ contributions

BI and MSI designed the study, MSI carried out the research under the supervision of BI. MSI carried out the statistical analysis and interpretation of data. MSI wrote the first draft. BI edited the final draft of the manuscripts. The authors read and approved the final manuscript.

## Funding

This work was partly supported by the Federal Government of Nigeria, through the Federal Scholarship Board under the Federal Ministry of Education Abuja, awarded to MSI (FSB/NA2020/1243), the Petroleum Technology Development Fund (Local Scholarship) awarded to MSI (PTDF/E/LSS/PG/ACI/Vol. 12/96) and the Admiralty University of Nigeria Staff development support, awarded to MSI (ADUN/REG/EGA/TS/8/35).

## Compliance with ethical standards

### Conflict of interest

The authors declare that no conflict exists among the authors.

## ADDENDUM

**Addendum 1:**
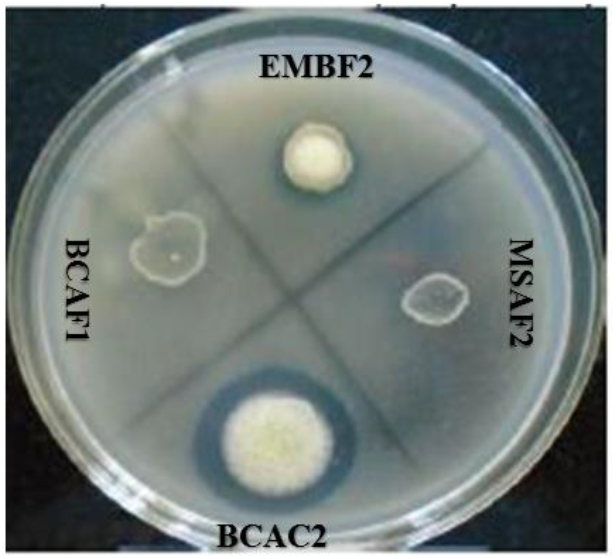
PSB showing efficiency in solubilization of insoluble phosphate by forming holo-zone [2].

**Addendum 3:**
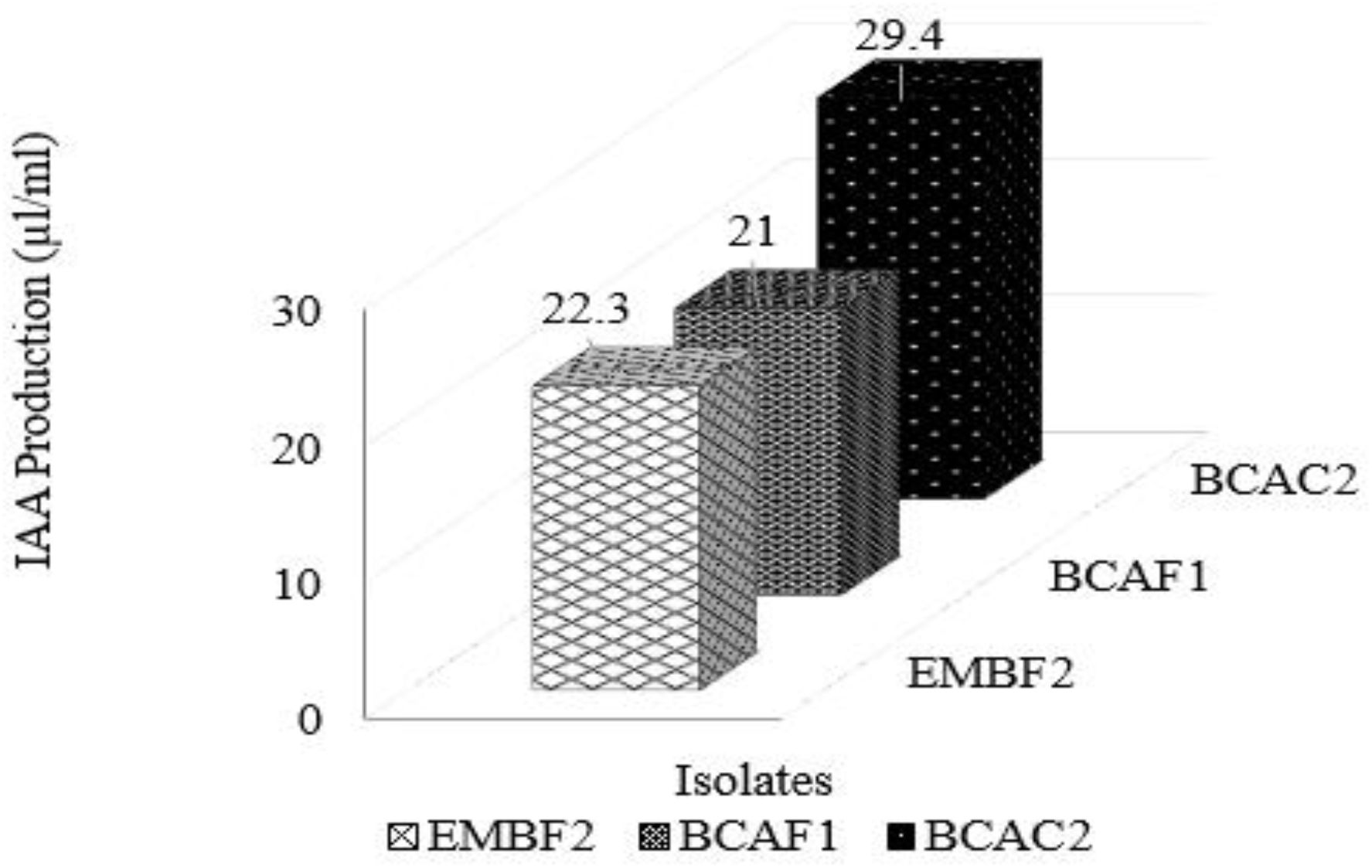
IAA production by the three isolated PSBs using Luria Bertani broth. BCAC2= *Bacillus cereus* strain GGBSU-1, BCAF1= *Proteus mirabilis* strain TL14-1, EMBF2= *Klebsiella variicola* strain AUH-KAM-9 [2].

**Addendum 4:**
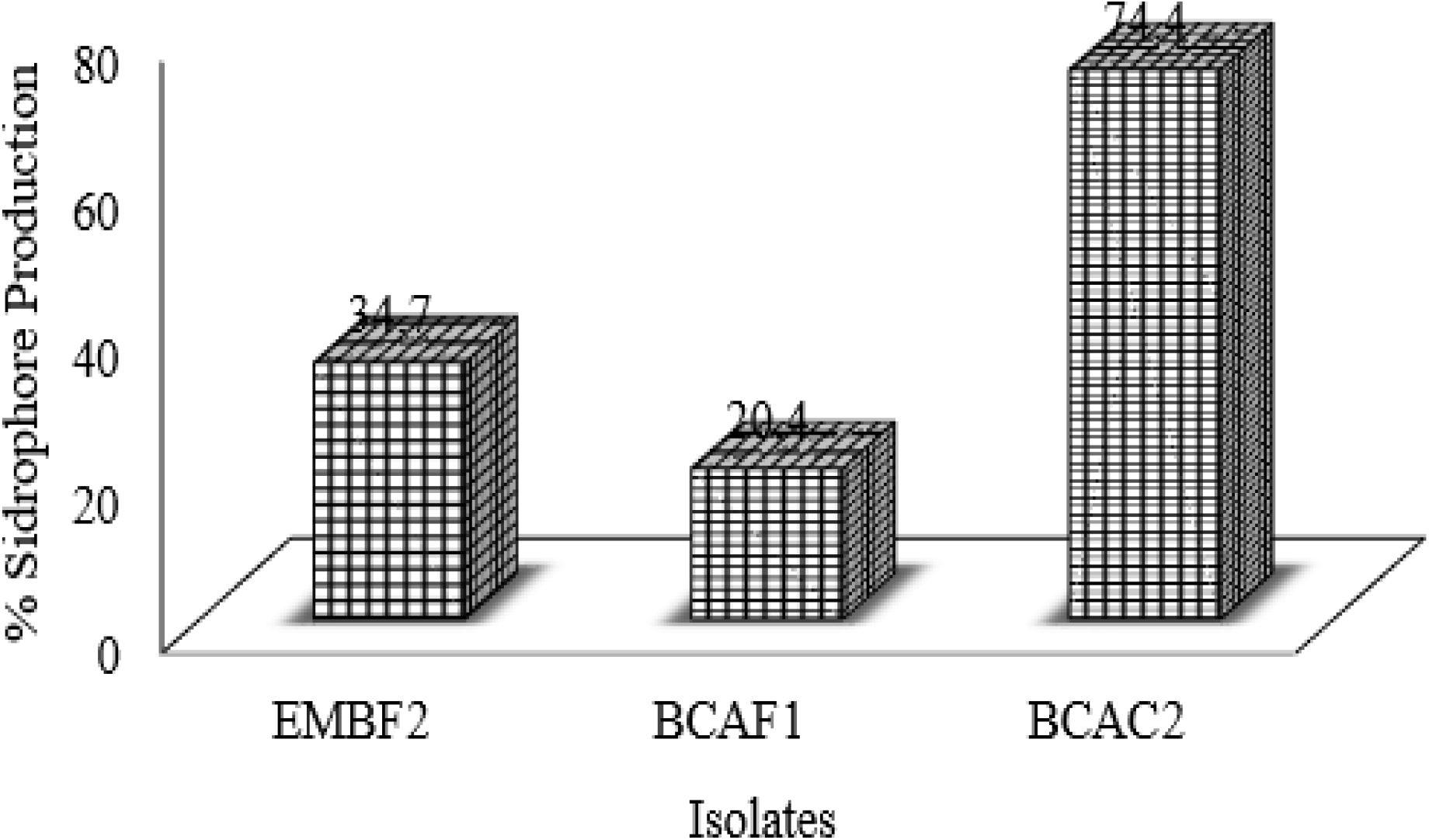
Percentage siderophores efficiency of the three isolated PSBs. BCAC2= *Bacillus cereus* strain GGBSU-1, BCAF1= *Proteus mirabilis* strain TL14-1, EMBF2= *Klebsiella variicola* strain AUH-KAM-9. [2].

**Addendum 5:**
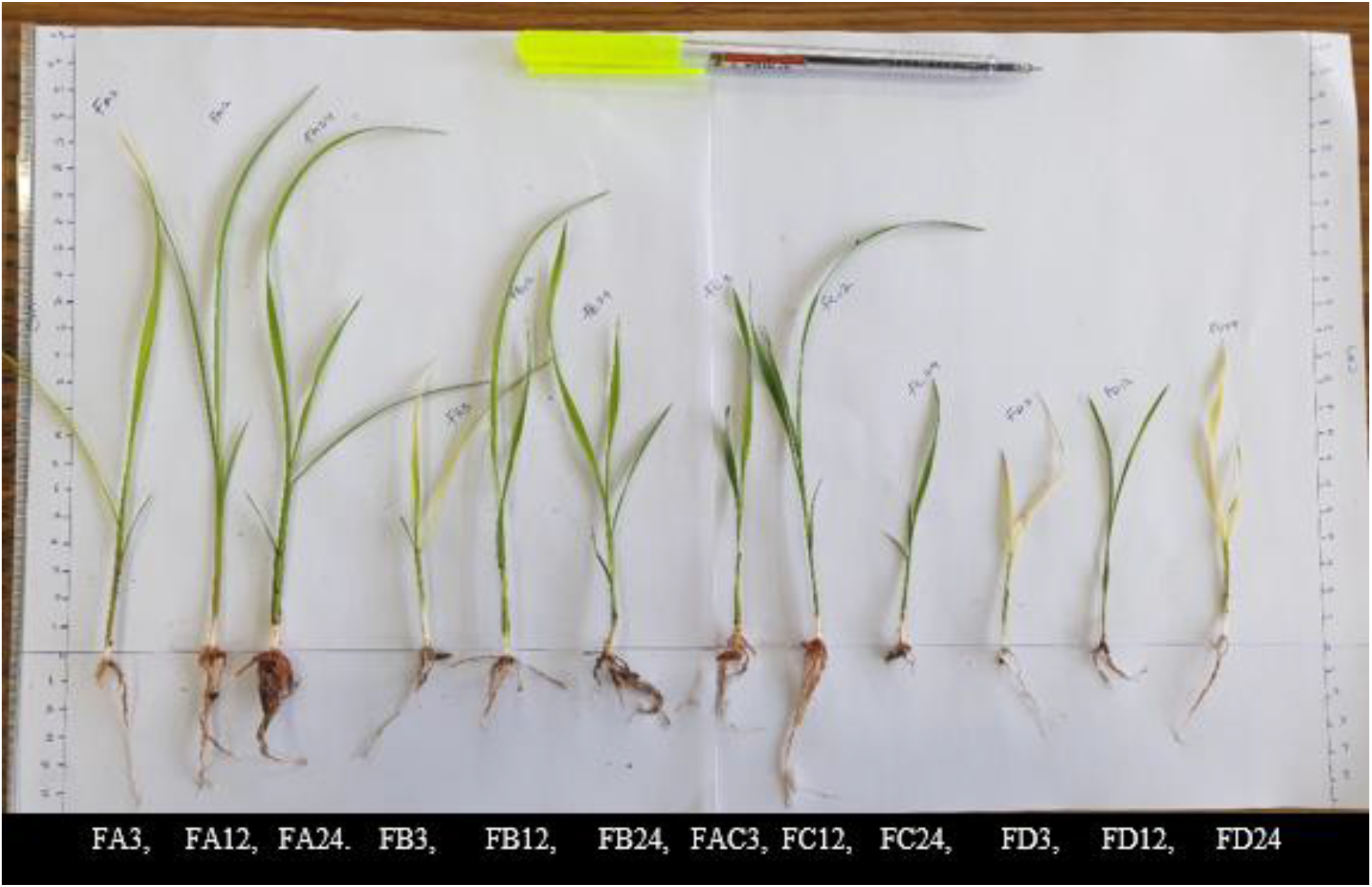
Morphological performance of rice seedling at 3 weeks after bio-priming.

## Notes

### Competing Interest Statement

The authors have declared no competing interest.

